# An Atlas of Short Linear Motif-Mediated Human Protein-Protein Interactions

**DOI:** 10.64898/2026.07.03.735260

**Authors:** Priyanka Madhu, Caroline Benz, Leandro Simonetti, Matthew J. Winters, Mythili S. Subbanna, Johanna Kliche, Lidia Gomez-Lucas, Hazem M Kotb, Izabella Krystkowiak, Dmitri Segal, William T. P. Darling, Sven Larsen-Ledet, Maximilian Vieler, Ana Zupancic, Aimiliani Konstantinou, Filip Mihalic, Julia K Varga, Livia Pagano, Susanne Lüchow, Andreas Kraemer, Lachlan Ellingboe, Kristof Görlitz, Germanna L. Righetto, Christin Kossmann, Ruisheng Xiong, Ora Schueler-Furman, Levon Halabelian, Vijayaratnam Santhakumar, Cheryl Arrowsmith, Stefan Knapp, Mate Erdelyi, Amelie Stein, Renaud Vincentelli, Peter M. Pryciak, Norman E Davey, Ylva Ivarsson

## Abstract

Short linear motifs (SLiMs) within intrinsically disordered protein regions mediate transient interactions crucial for cell physiology^1^. However, the global interaction landscape of human SLiMs remains largely uncharted. Here we present the Atlas of SLiM-mediated Human protein-protein Interactions (ASHI), which maps more than 20,000 interactions by screening over 800 human protein domains against a library of one million peptides tiling the human disordered proteome. ASHI expands the SLiM interactome, uncovers novel binding modes for known peptide-binding domains, and reveals unexpected peptide-binding activities in enzymes, chaperones, RNA-binding proteins, and modification-reader domains. Furthermore, intrinsically disordered regions emerge as densely encoded interaction platforms where interaction specificity is governed by diverse mechanisms, including key motif determinants, flanking residues, competition, and multivalency. These data provide an unprecedented foundation for modeling dynamic interaction networks, interpreting disease-associated variants, and decoding the dark proteome.

## Introduction

Cellular behavior depends on a dense and dynamic network of protein-protein interactions that coordinate signaling, gene expression, and proteostasis. Our understanding of the human interactome has been greatly expanded by proteome-scale approaches^2–5^, which have systematically revealed protein localization patterns, complex assemblies, and direct binary interactions^3^. More recently, large-scale structure prediction has provided new insights into the organization and architecture of macromolecular assemblies^6,7^. Among the most elusive components of the interactome are short linear motifs (SLiMs): short (typically 3–10 amino acids), degenerate peptide sequences embedded within intrinsically disordered protein regions (IDRs), that are recognized by folded peptide-binding domains in their interaction partners^8^. SLiMs are important functional elements of IDRs and provide sites for scaffolding, enzyme docking, cargo selection, and spatial targeting^8^. Although short and sequence-degenerate, SLiMs achieve high-fidelity cellular selectivity. The mechanisms by which these seemingly low-information motifs drive precise recognition remains unresolved at the proteome scale. Strikingly, dysregulation of SLiM-mediated interactions is increasingly linked to human disease. Viruses exploit host pathways through molecular mimicry of SLiMs^9^, and disease-associated mutations in IDRs often disrupt or create SLiM–domain interactions^10^, underscoring the therapeutic importance of charting the SLiM-mediated interactome.

Although the low to moderate affinity of SLiMs enables rapid, reversible network rewiring, it also makes them exceptionally difficult to capture experimentally, which creates a major blind spot in the human interactome^11^. While bioinformatic analyses estimate the human proteome may contain over 100,000 SLiMs^12^, only a small fraction are currently catalogued at the Eukaryotic Linear Motif (ELM) database (2,451 instances)^8^ or the Motif Map of the Proteome (MoMaP) database (7,119 instances; https://slim-tools.org/momap/), suggesting that most motif-mediated interactions remain to be discovered. Recent methodological advances have enabled direct investigation of this unexplored interaction space. Peptide-discovery platforms such as phage, yeast, and mRNA display have transformed our ability to interrogate peptide recognition at scale^13^. We established a proteomic peptide-phage display (ProP-PD) approach for systematically charting the SLiM-based interaction space^14^. A second-generation human disorderome (HD2) library encodes one million overlapping peptides that provide dense coverage of all human IDRs (**Fig. 1a**)^14^. A pilot screen of this library against 35 bait domains yielded over 2,000 interaction pairs^14^, demonstrating feasibility while underscoring the vast, unmapped landscape of motif-mediated biology.

**Figure 1.**
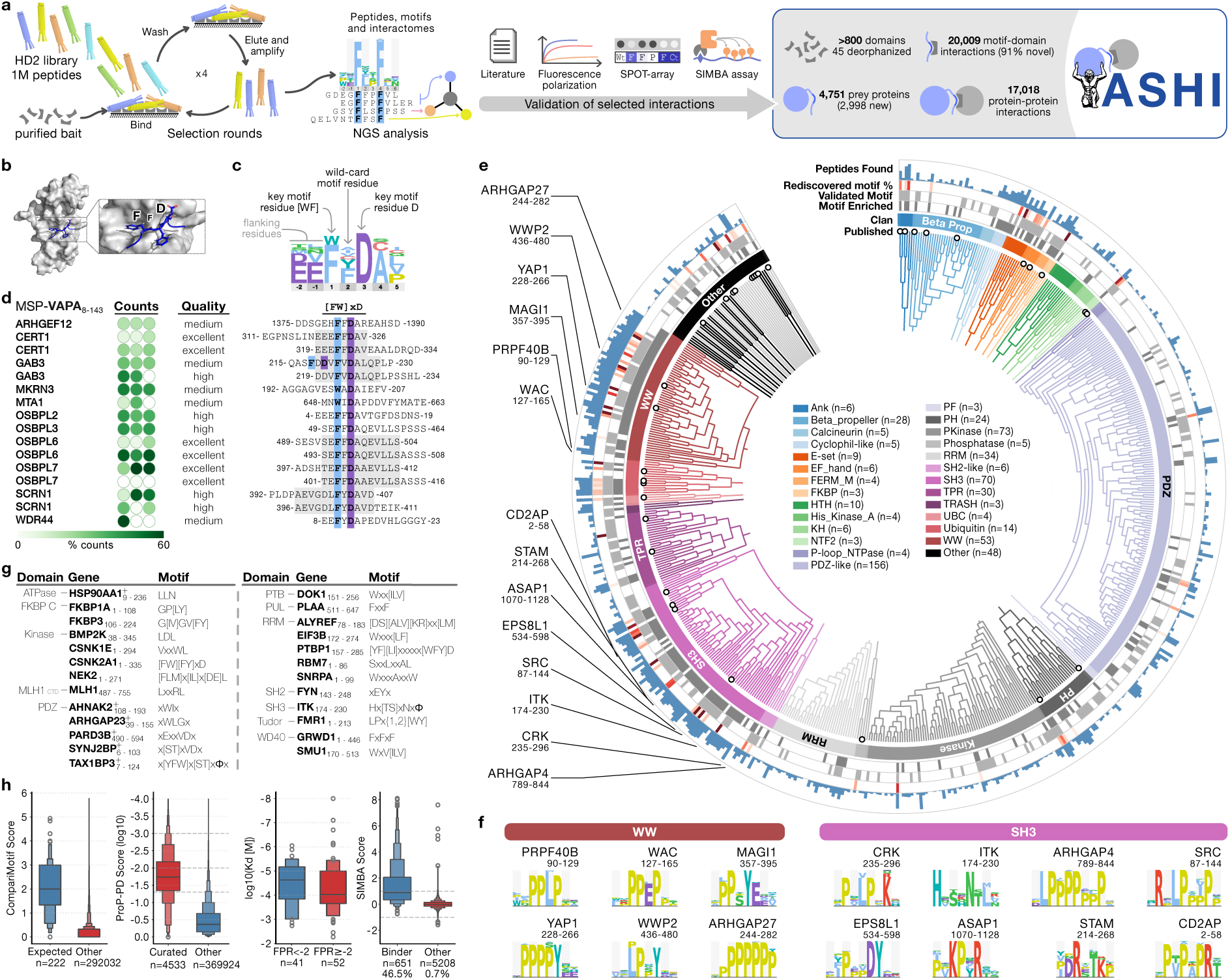
The Atlas of SLiM-mediated Human protein-protein Interactions (ASHI). **a,** Schematic overview of the generation of ASHI, starting from ProP-PD selections (left) using the HD2 that tiles the human IDRs, data analysis and orthogonal validations (middle) and the resulting ASHI. **b–d,** Representation of a SLiM-mediated interaction and generated results using the VAPA MSP domain as a representative example. **b**, Example of a VAPA MSP domain bound to a [FW]xD motif in the oxysterol-binding protein 1 (OSBP1 357-363; PDBid 2RR3). **c**, Definition of key, wildcard and flanking residues within VAPA MSP binding peptides, as shown using a position specific scoring matrix (PSSM) representation of the motif. **d**, Representative example of the information generated by the ProP-PD screening, including enrichment (% of NGS counts in each of 3 replicates), quality of data (medium to excellent), peptides and motifs (the shaded part of the sequences indicate overlap between identified adjacent peptide tiles; each peptide in the library is 16 amino acid long, and the overlap between tiles are 12 amino acids). **e**, Overview of the results provided in ASHI (number of peptides found/bait; rediscovery of known motifs; enriched motifs). **f**, Representative motifs identified for the WW and SH3 domain families. **g**, Overview of newly defined motifs in ASHI. The “+” mark next to gene names indicates that the motif was found also for other domain family members and a representative example is shown. **h,** Quality assessment of the ASHI dataset. Left, comparison of motif recovery for previously reported motifs of the screened baits (blue) relative to all other motifs. Middle left, comparison of ProP-PD confidence scores for previously curated binding peptides and other peptides. Middle right, K_d_ values measured by FP for high-confidence versus lower-confidence ligands. Right, SIMBA scores for bait-specific peptides compared with non-cognate peptide–domain pairs.

In this study, we present ASHI (the Atlas of SLiM-mediated Human protein-protein Interactions), a resource combining proteome-scale experimental mapping with high-throughput affinity ranking to screen more than 800 purified human protein domains against the HD2 library. This effort yields over 20,000 domain-motif interactions, more than doubling the known motif-based interactome, and reveals that SLiM recognition is organized across multiple scales. Globally, we show that peptide recognition is a far more widespread property of the proteome than currently appreciated, extending to canonical RNA-binding domains, post-translational modification readers, and major chaperones. At the motif level, context tunes domain specificity beyond the core consensus. At the level of full IDRs, overlapping, repeated and co-occurring motifs create opportunities for competition, avidity and multi-partner recruitment. Finally, at the network level, SLiM-containing proteins are themselves embedded in interconnected assemblies that include SLiM-binding proteins, essential proteins, disease-associated variants and therapeutic targets. Together, ASHI provides a framework for decoding a layered interaction logic of disordered human proteome.

## Results

### A proteome-scale SLiM-based interaction atlas

To systematically map SLiM-based interactions across the human proteome, we screened over 800 purified human protein domains, spanning major peptide-binding families and a broad range of less-characterized folds (**Supplementary Table 1**). The domain set was screened against the HD2 ProP-PD library, a million-peptide phage display library tiling all human IDRs^14^ (**Fig. 1a-d**). This library was specifically designed to capture motif-containing peptide sequences embedded within IDRs that bind to folded domain baits. Enriched phage pools were analyzed by next-generation sequencing (NGS), identifying peptide ligands for 525 domains present in 430 proteins (**Supplementary Table 2**). Integrative scoring based on reproducibility, peptide overlap, NGS enrichment, bait specificity, and consensus similarity (**Fig. 1d; Supplementary Methods; Extended Data Fig. 1**) identified 20,009 motif-domain interactions (17,173 if excluding peptides predicted to be in structured regions according to AlphaFold3^15^). For consistency, previously released ProP-PD data^14,16–18^ was reprocessed, complementing the ASHI data set with 2,564 motif-motif interactions (**Supplementary Table 2**). Cross-referencing against the MoMaP database (April 2026), 320 of the 3,055 motif instances previously reported to bind the screened domains were rediscovered (**Supplementary Table 3**). The motifs were found in 4,751 prey proteins, of which 2,998 (63%) had no prior annotation as being SLiM containing. The screen also deorphanized 45 domains by revealing their peptide ligands for the first time (**Supplementary Table 2)**, establishing SLiM-based recognition as a more widespread feature of the human proteome than previously observed.

Our screen revealed distinct dimensions of novelty. First, we identified motif-binding capacity in domains that have not historically been recognized as peptide-binding modules, including RNA-processing regulators, isomerases, and chaperones (**Fig. 1e**). Second, several established motif-binding domain families recognize motif signatures diverging substantially from their expected consensus (**Fig. 1g**), pointing toward structural plasticity or alternative binding surfaces as mechanistic explanations. Thus, even well-studied folds showed unrecognized motif-binding capabilities. Crucially, the most baits recapitulated their expected motif consensus (**Fig. 1f, h; Extended Data Fig. 2**), validating the robustness of our screening platform.

We validated representative interactions using orthogonal binding assays. Affinities were determined using fluorescence polarization (FP) competition assays for 29 domains and using ligands spanning the full ProP-PD confidence range (**Fig. 1h; Supplementary Table 4**; **Extended Data Fig. 3**). Of the 103 interactions tested, 93% (96) displayed measurable binding, with K_d_ values spanning the nanomolar to millimolar range (median: 30 µM). Higher ProP-PD scores were associated with significantly lower K_d_ values (**Fig. 1h**), supporting a correspondence between screen enrichment and affinity. To extend affinity ranking, we employed Systematic Intracellular Motif Binding Analysis (SIMBA)^19^, a quantitative yeast-based peptide-domain binding assay. Notably, we observed a clear correlation between SIMBA scores and the FP-determined affinities (**Extended Data Fig. 4a-i**). Of 649 tested interaction pairs across nine domains, 46.2% (300) were confirmed binders with a SIMBA score >1 (**Fig. 1h; Supplementary Table 5, 6**), with a 67.1% validation rate among the higher confidence ProP-PD ligands (**Extended Data Fig. 4j)**. Only 0.7% of the expected non-binding pairs interacted according to the SIMBA assay. The results demonstrate that ProP-PD screens effectively identified genuine biophysical interactions.

Collectively, ASHI more than doubles the catalog of SLiM-based human interactions relative to the MoMaP database (April 2026) and establishes a proteome-scale framework to decode the molecular rules governing motif recognition. Focusing on the key findings, the following sections evaluate novel ligand recognition by canonical motif-binding domains, identify unexpected peptide binding events that represent novel motif classes, and highlight broader principles of motif biology uncovered by our platform.

### Expanding the interactomes of canonical SLIM-binding domains

Several large domain families are canonically associated with motif binding, mediating numerous protein–protein interactions that underpin eukaryotic signaling networks. Two of the best-characterized examples are the Src Homology 3 (SH3) and WW domain families, which recognize proline-rich PxxP and [LP]PxY motifs respectively. While previous studies have defined broad binding consensus motifs and numerous partners^20,21^, proteome-wide mapping of their binding sites in human proteins at motif-level resolution has remained lacking. To address this, we applied ProP-PD to 62 SH3 and 57 WW domains, enabling proteome-scale interrogation of their interaction landscapes (**Fig. 1e,f**). We recaptured previously defined SH3/WW binding preferences, including classical PxxP/[LP]PxY motifs as well as known non-canonical specificities (**Fig. 1f**). A notable outlier was the ITK SH3 domain, which exhibited dual specificity, enriching both the canonical PxxP motif and a distinct non-canonical HxSxNxL consensus. Beyond validating specificity determinants, we rediscovered known binding motif instances (SH3 domains: 75 of 399; WW domains: 118 of 221) while substantially expanding the number of interactions by identifying 2,381 and 1,355 additional binding sites for SH3 and WW domains, respectively (**Supplementary Table 2**). This increased coverage is particularly impactful for understudied family members, and 45 of these domains were effectively deorphanized in terms of ligand binding. Collectively, these data establish an atlas of SH3 and WW binding motifs across the human proteome.

We next extended our approach to peptide binding domain families with more limited prior knowledge on SLiM binding. WD40 domains are β-propeller folds that serve as versatile interaction scaffolds capable of recognizing a broad range of ligands^22^. Many WD40 family members have no known ligands, motivating us to apply ProP-PD to 19 WD40 domains. Two of these, SMU1 DNA replication regulator and spliceosomal factor (SMU1) and glutamate rich WD repeat containing 1 (GRWD1), showed specific and reproducible binding to peptides (**Supplementary Table 2**). For SMU1, which is involved in pre-mRNA splicing^23^, a WxV[ILV] consensus motif was defined. A SMU1 binding peptide was found in the 7S U2 SnRNP spliceosome complex component HTATSF1; this ligand was modelled by AlphaFold3^15^ to bind to the rim of the domain (**Extended Data Fig. 5a**). The WD40 domain of GRWD1, a histone binding protein^24^ also implicated in ribosomal biogenesis^25^, recognized instead a distinct FxFxF motif, for which structural modeling proved less conclusive. Nevertheless, two tested GRWD1-interacting peptides bind with high affinity (K_d_ values C8orf33: 3.0 µM and SELENOI: 4.2 µM; **Supplementary Table 4, Extended Data Fig. 3g, 5b**). A highly conserved FxFxF motif is present in a reported GRWD1 interactor, histidine protein methyltransferase 1 homolog (METTL18)^26^, supporting a potential biological relevance of the defined specificity.

Together, these findings demonstrate how systematic ProP-PD screening simultaneously broadens the interaction landscapes of canonical motif-binding families and deorphanizes their unstudied members.

### SLiM binding by enzymes and chaperones

Beyond canonical motif-binding domains, the screen revealed SLiM-based recognition across a broad range of enzymatic and chaperone domains. Identified ligands engage diverse binding surfaces, including kinase docking grooves, auxiliary surfaces in molecular chaperones and catalytic pockets of peptidyl–prolyl isomerases (PPIases).

### Kinase domains recognize diverse motifs

Kinases exploit SLiM-based interactions for substrate targeting, regulatory control, and complex assembly^27^. In tyrosine kinases, these interactions are commonly mediated by auxiliary SH2 and SH3 domains, and our ProP-PD screen yielded many SH3-mediated interactions across non-receptor tyrosine kinases (**Supplementary Table 2**). Serine/Threonine kinases employ more diverse mechanisms, including auxiliary domains, regulatory subunits, and direct docking of SLiMs to the kinase domain at sites distinct from the catalytic cleft. Screening of 96 kinase domains identified peptide ligands for 63 kinases, revealing both known and novel consensus motifs. For MAPK family members, ligands were found with D-site-like motifs^28^, as well as a novel ExRxxLxL motif in WDR62 that bound MAPK8 (JNK1) with nanomolar affinity (**Fig. 2a**). CASK recognized instead a x[IV]Wx consensus motif that bound with low-micromolar affinity (**Fig. 2b, f**) and which AlphaFold3^15^ modeling predicted to bind a surface partially overlapping the D-site docking pocket of MAPKs (**Fig. 2k**). CSNK1D and CSNK1E ligands contained a [IV]xxWL motif (**Fig. 2f; Extended Data Fig. 2**), which was validated by alanine scanning peptide array (**Fig. 2g**) and was modeled to a surface on the kinase C-lobe domain (**Fig. 2k**). NEK2 peptide ligands contained a [FLM]x[IL]x[DE]L motif, which bound with mid-micromolar affinities and was validated by alanine scanning (**Fig. 2d, h**). A representative RSF1 peptide was modeled with AlphaFold3 to interact with the NEK2 N-lobe (**Fig. 2j**) analogous to the TPX2-binding surface on Aurora A^29^. Consistent with biological relevance, the NEK2-binding motif in NEK11 overlaps the region previously implicated in the NEK2-NEK11 interaction^30^. For Aurora A, we identified a TROAP-derived peptide (**Fig. 2e**) with similar sequence as the known Aurora A peptide ligand from TACC3^31^, modelled to bind a distinct surface on the N-lobe (**Fig. 2j**). These data collectively demonstrate that kinases interact with a variety of SLiMs using distinct interfaces.

**Figure 2.**
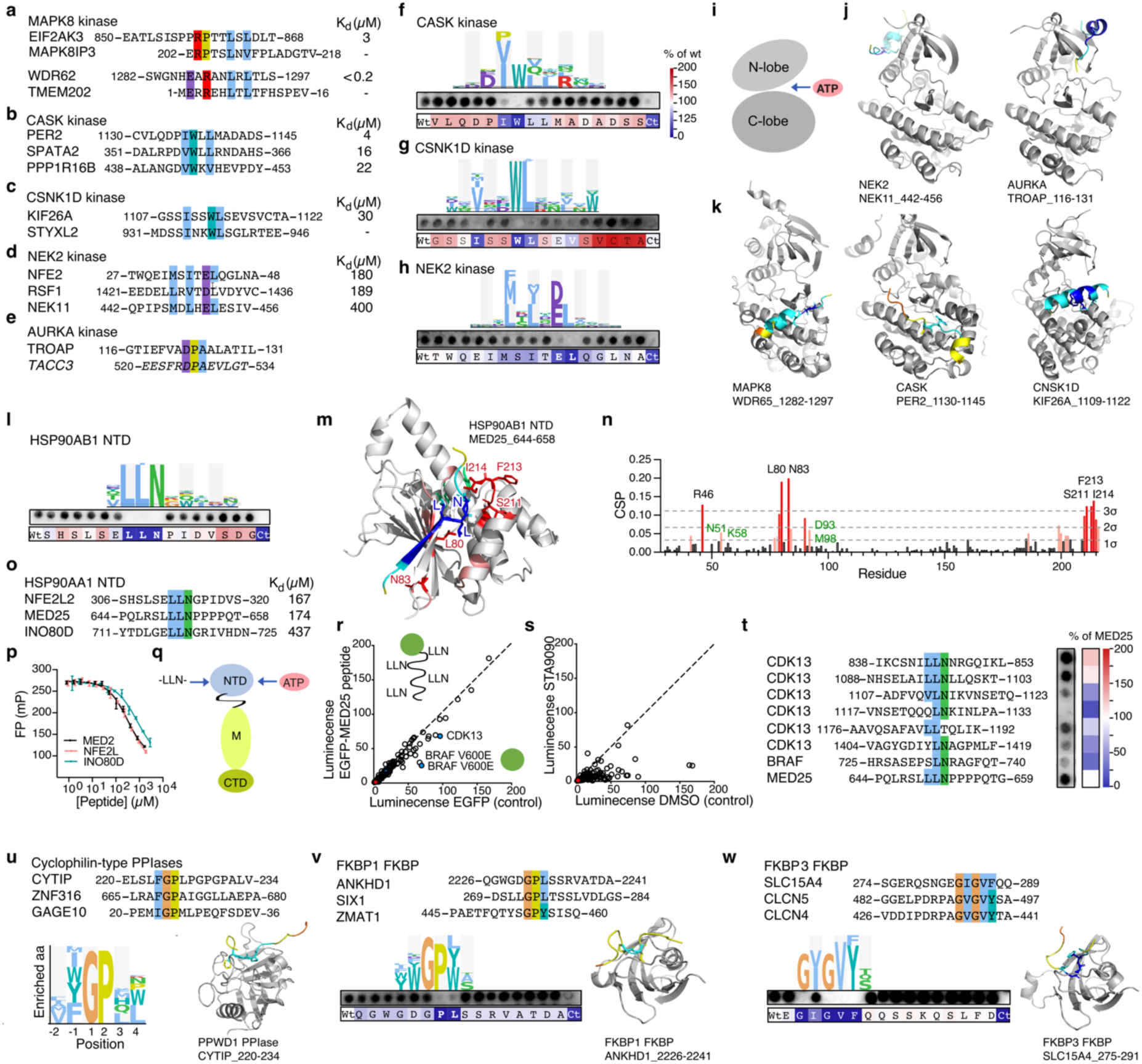
SLiM binding by enzymes and chaperones supports substrate recognition, complex assembly and regulation. **a–e,** Selected ligands of MAPK8 (**a**), CASK (**b**), CSNK1D (**c**), NEK2 (**d**) and AURKA (**e**) kinase domains, with corresponding dissociation constants (K_d_) measured by competitive FP (n=3). **f–h**, PSSMs for motifs defined for CASK (**f**), CSNK1D (**g**) and NEK2 (**h**) kinase domains, together with alanine-scanning peptide arrays of representative peptide ligands from PER2 (**f**), KIF26A (**g**) and NFE2 (**h**) validating core motif residues. Spot intensities (n=2) are shown as percentage of wild-type binding and range from 200% to 0%, as indicated in (**f)**. **i**, Schematic of a kinase domain showing the N-lobe, C-lobe and ATP-binding site. **j,k**, AF3 models of peptide binding to distinct sites on the N-lobes of NEK2 (right; ipTM 0.78, pTM 0,86) and AURKA (left; ipTM 0.86, pTM 0.86) (**j**) and on the C-lobes of MAPK8 (right; ipTM 0.82, pTM 0.86), CASK (middle; ipTM = 0.68p, pTM = 0.86) and CSNK1D (left)(**k**). **l**, PSSM of the HSP90AB1 N-terminal domain (NTD)-binding motif (top) and alanine-scanning peptide array validating the motif (bottom). Spot intensities are shown as in **f**. **m,n**, NMR chemical-shift perturbations mapped onto an AF3 model of the MED25 peptide–HSP90AB1 NTD complex (ipTM 0.78, pTM = 0.87). In (**m)**, the peptide is colored by AF3 confidence and HSP90AB1 NTD is shown in grey. In **n**, perturbations observed upon addition of five molar equivalents of peptide are plotted by residue; residues exceeding 1, 2 or 3 standard deviations are indicated in black, light red and red, respectively. Residues with the strongest perturbations are highlighted in (**m)**, and ATP-contacting residues are shown in green. The proposed peptide-binding site is opposite the ATP-binding pocket. **o-p**, Affinities of HSP90 NTD-binding peptides measured by competitive FP using geldanamycin as probe (n=3). **q**, Schematic of the relative positions of the ATP-binding site and peptide-binding site within the HSP90 NTD in the context of the middle (M) and C-terminal (C) domains. **r,s**, Normalized luminescence values from LUMIER interaction assays showing the effects of an EGFP-tagged MED25 peptide containing four motif repeats (**r**) or the HSP90 ATP-pocket inhibitor STA9090 (**s**) on HSP90 interactions with indicated client proteins. The MED25 peptide selectively competed interactions with CDK13 and BRAF V600E, whereas STA9090 had broader inhibitory effects (n=2). **t**, Peptide-array analysis of predicted HSP90 NTD-binding motifs in CDK13 and BRAF. Four motifs were validated in CDK13, whereas a putative motif in BRAF showed weak binding. **u**, Representative ProP-PD ligands of cyclophilin-type peptidyl–prolyl isomerases (PPIA, PPIB, PPIE, PPWD1 and NKTR) aligned to show a shared motif (top), with a representative position-specific scoring matrix (PSSM; left) and an AlphaFold3 model of the PPWD1 isomerase domain bound to the CYTIP_220–234_ peptide (right; ipTM = 0.65, pTM = 0.91). **v**, Representative ligands of the FKBP1A FKBP domain with a shared GP[LY] motif (top), alanine-scanning peptide array of the ANKHD1_2226–2241_ peptide validating the PL core motif (left), and AF3 model of the complex (right; ipTM = 0.52, pTM = 0.85). Spot intensities are shown as percentage of wild-type binding, from white (100%) to dark blue (0%). **w**, Representative ligands of the FKBP3 FKBP domain with a shared G[IV]GV[FY] motif (top), alanine-scanning peptide array of the SLC15A4_282–297_ peptide validating the GxGVF motif (left), and AF3 model of the complex (right) (n=2). See **Supplementary Figure 4c-k** for alternative representations of the modelled complexes.

### SLiM binding by the N-terminal domain of the chaperone HSP90

We next examined peptide binding by three regions of the molecular chaperones HSP90α and HSP90β: the N-terminal ATPase domain (NTD), the middle domain and the C-terminal domain. We uncovered peptide-binding to the NTDs and defined an LLN consensus motif (**Fig. 2l**). The set of LLN motif containing proteins is enriched in proteins linked to transcriptional regulation (**Supplementary Table 2**), including the Mediator of RNA polymerase II transcription subunit 25 (MED25) and the nuclear factor erythroid 2-related factor 2 (NFE2L2). Alanine-scanning peptide arrays confirmed the LLN consensus (**Fig. 2l**), and NMR chemical-shift mapping localized the binding site of the MED25 peptide to a HSP90 NTD surface pocket distinct from the ATP-binding cleft (**Fig. 2m,n; Supplementary Table 7**). The LLN-binding surface overlaps with an intramolecular HSP90 contact involving an ELN segment (residues 289–291) at the end of the flexible NTD–middle-domain linker based on available structures^32^.

The chemical-shift perturbations propagate toward the nucleotide-binding site (**Fig. 2m,n**), suggesting communication between the peptide-binding surface and the ATPase pocket. Consistent with such a coupling, competitive FP based affinity measurements using FITC-labeled geldanamycin (which binds to the ATP site) showed that LLN-containing peptides displaced geldanamycin with mid-micromolar affinities (**Fig. 2o-q)**. Direct binding with FITC-labeled peptides confirmed the observed affinity range (**Supplementary Table 4**). In cells, LUMIER-based competition experiments using a GFP-tagged MED25 peptide showed that expression of the LLN motif selectively displaced a subset of HSP90 clients, whereas addition of the HSP90 NTD inhibitor STA9090 affected a broader set of clients (**Fig. 2r,s**). The LLN peptide affected HSP90 binding of CDK13 and BRAF^V600E^ but not wild-type BRAF. This is notable as BRAF^V600E^, but not BRAF wild-type, is dependent on the HSP90-CDC37 complex for stability and function^33^. Sequence inspection of BRAF revealed that the protein has a partially matching “LN” motif which showed weak but detectable binding (**Fig. 2t**), although this does not explain the differential effect observed for wild-type and mutant BRAF. CDK13 contains two LLN motifs, as well as additional partial matches, which bind readily to HSP90 (**Fig. 2t**). These findings identify the HSP90 NTD as a peptide-recognition module and suggest that a subset of chaperone–client interactions can be modulated by SLiM docking to a regulatory NTD surface.

### Substrate and ligand binding by the catalytic domains of proline isomerases

PPIases catalyze cis–trans isomerization of proline-containing peptide bonds, thereby governing protein folding and conformational dynamics^34^. In our screens, members of the cyclophilin family (NKTR, PPIA, PPIB, PPIE, PPWD1) enriched partially overlapping peptide sets (**Supplementary Table 2)** containing [FY]GP-related motifs. These motifs were modelled onto the active site and resemble known cyclophilin substrates^35^ (**Fig. 2u**), and the results are in line with recent evidence that cyclophilins facilitate translation of intrinsically disordered proteins^36^. The structurally distinct PPIase FKBP1A recognized GP[LY]-containing motifs (**Fig. 2v)**, whereas the isomerase domain of FKBP3 bound non-proline-containing peptides (**Fig. 2w**), indicating a distinct ligand tolerance. The FKBP3 ligands shared a G[IV]GV[FY] stretch, and the importance of each position was confirmed by alanine scanning peptide array. This suggests that some PPIase domains, such as FKBP3, may serve dual functions as isomerases and as SLiM-binding modules recognizing distinct sequences.

### Unexpected peptide binding by well-studied domains

The ASHI dataset further uncovered non-canonical modalities of SLiM-based interactions, including motif engagement by classically post-translational modifications (PTMs)-dependent domains, PDZ domains binding to internal sequences, and SLiM recognition by RNA binding domains.

### PTM-reader domains also recognize unmodified peptide ligands

PTM readers are traditionally classified by their strict dependency on chemical modifications. For instance, SH2 and PTB domains are prototypic readers of phospho-tyrosine containing peptides^37^. Intriguingly, we found that a subset of these domains could bind to unmodified ligands, including the SH2 domain from FYN and the PTB domain of DOK1. For the FYN SH2 domain, we uncovered interactions with peptides from POC5 and PRUNE2 (**Fig. 3a**). Unphosphorylated PRUNE2 bound FYN SH2 with low but measurable affinity (217 µM K_d_), and phosphorylation increased the affinity (3.7 µM K_d_). In comparison, a high affinity viral phosphopeptide (from the Middle T antigen, (POVHA)^38^) binds with sub-micromolar affinity (**Fig. 3a**). Peptide array alanine scanning revealed that the binding of the PRUNE2 peptide involves an IPEY stretch plus an additional distal glutamic acid (**Fig. 3a)**. The IRS type PTB domain of DOK1 bound unmodified ligands harboring a Wxx[ILV] motif, including a PTPN23 derived peptide which docked to part of the PTB binding groove (**Extended Data Fig. 5l**). This PTPN23 peptide bound with lower affinity than the unphosphorylated or phosphorylated peptides from the known DOK1 ligand RET^39^ (**Fig. 3b**). Notably, the PTPN23 peptide lacks a tyrosine, usually present in PTB binding peptides. Alanine scanning (**Fig. 3b**) established that the PTPN23 motif is distinct from the canonical PTB binding NPxpY motif and instead resembles a previously described talin (TLN1) PTB binding motif^40^, which is anchored by a tryptophan residue and shows permissiveness to motif variations based on DMS analysis (**Extended Data Fig. 6a,b**). Binding of tryptophan-anchored motifs thereby emerges as a shared feature among the IRS-type PTB domains.

**Figure 3.**
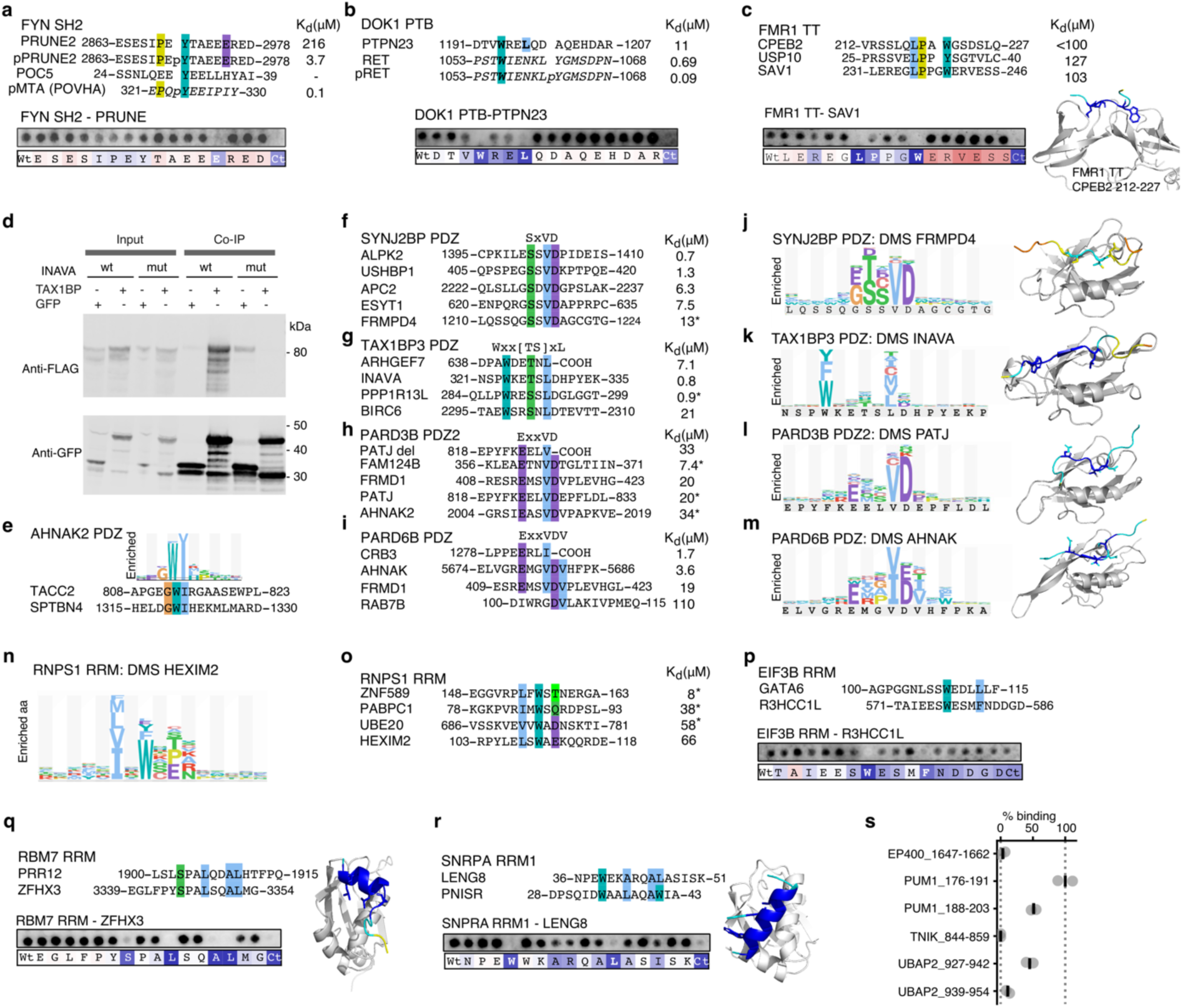
Unexpected modes of SLiM-binding: Internal PDZ binding motifs, SLiM-binding RRM domains and PTM binders that bind unmodified ligands. **a**, FYN SH2-domain ligands from ProP-PD, affinities of unmodified and phosphorylated PRUNE2 peptides as determined by FP (n=3), and alanine-scanning of the PRUNE2 peptide (n=2). A high-affinity viral phospholigand (pMTA) is shown for comparison. **b,** Comparison of DOK1 PTB-domain ligands from PTPN23 and RET, with affinities (n=3) and alanine-scanning validation of the DOK1 PTB binding motif in the PTPN23 peptide (n=2). **c,** FMR1 tandem Tudor (TT) domain ligands, affinities (n=3), alanine-scanning validation (n=2) and AF3 model of the FMR1 TT–CPEB2_216-224_ complex (ipTM = 0.83, pTM = 0.86). **d**, Co-immunoprecipitation of full-length FLAG-tagged INAVA with GFP-tagged TAX1BP3 wild-type or motif mutant (324-WKETSLD-330 mutated to 324-AKEAAAA-330) expressed in HEK293 cells (n=3). **e**, PSSM of AHNAK2 PDZ-binding peptides and alignment of two representative ligands.**f–i**, Representative ligands of SYNJ2BP PDZ (**f**), TAX1BP3 PDZ (**g**), PARD3B PDZ2 (**h**) and PARD6B PDZ (**i**), with affinities determined by competitive FP (n=3) or estimated from SIMBA scores (*; n=6). Affinities of C-terminal ligands are shown for comparison. **j–m**, DMS-derived PSSMs and AF3 models for selected ligands of SYNJ2BP PDZ (j; pTM 0.42, pTM 0.8), TAX1BP3 PDZ (**k**; ipTM = 0.77, pTM = 0.83), PARD3B PDZ2 (**l**; ipTM 0.77, pTM 0.85) and PARD6B PDZ (**m**), together with AF3 models of the complexes. **n,o**, RNPS1 RRM peptide binding. DMS of a HEXIM-derived peptide confirming a hydrophobic motif (**n**) together with estimated affinities by FP-based affinity measurement (HEXIM2; n=3) or SIMBA based affinity estimates (*; n=6) (**o**).**p–r**, RRM-binding peptides of EIF3B (**p**), RBM7 (**q**) and SNRPA (**r**) with alanine-scanning peptide array validation (n=2) and AF3 models (RBM7-ZFHX3: ipTM 0.79, pTM 0.86; SNRPA RRM1 – LENG8: ipTM 0.85, pTM 0.82). **s**, Peptide SPOT array analysis of a set of potential PCBP1 KH1 ligands (n=2). See **Supplementary Figure 4k-r** for additional representations of the modelled complexes.

Together, the SH2 and PTB results highlight that, while the high affinity ligands are phosphorylated, these domains can exhibit latent modification-independent specificity. This phenomenon extends beyond phosphorylation readers to other classes of modification-dependent modules, such as the tandem Tudor (TT) domain of the Fragile X messenger ribonucleoprotein 1 (FMR1), which typically binds methylated residues^41^. In our screens, this domain enriched peptides with a previously undescribed LPx{1,2}[WFY] consensus motif, validated by alanine-scanning peptide arrays (**Fig. 3c**). AlphaFold3 modeling of a CPEB2-derived peptide (**Fig. 3c**) suggested that the tryptophan side chain inserts into the aromatic cage typically occupied by methylated lysine residues^42^. These observations support that some PTM-binding domains possess broader interaction capacities than just their canonical ligands.

To evaluate the systemic interplay between modifications and motif recognition at a global scale, we examined the PTM status across the domain-binding IDRs. Of the domain-binding IDRs, 2,825 contain annotated PTM sites (**Supplementary Table 2**), which potentially may affect roughly one-third of all interactions reported. As expected by its abundance, phosphorylation was by far the most prevalent modification and this PTM is likely to tune the affinity of numerous of the SLiM-based interactions (**Extended Data Fig. 7**).

### Internal-motif recognition is widespread across the human PDZ domain family

While the textbook definition of PDZ domains dictates a strict requirement for terminal carboxyl groups^43^, our systematic screening reveals that this positional constraint is vastly oversimplified. ProP-PD profiling of all 266 human PDZ domains revealed that more than 40% (118/266) engage internal peptide motifs (**Supplementary Table 2)**. Supporting the functional relevance of these non-canonical interactions, co-immunoprecipitation confirmed the interaction between Tax1-binding protein 3 (TAX1BP3) and the internal PDZ binding motifs in the innate immunity activator protein (INAVA) (**Fig. 3d**). SIMBA-based affinity ranking combined with FP-monitored measurements confirmed multiple low-micromolar interactions across diverse PDZ domains, including SYNJ2BP, PARD3B PDZ2, PARD6B, and TAX1BP3 (**Fig. 3f-i**). The affinities of non-canonical internal PDZ ligands were comparable to those of the C-terminal ligands tested here (**Fig. 3f-i**) and reported elsewhere^43,44^. The internal PDZ ligands commonly retain canonical recognition features despite lacking a free C-terminus, with a subset bearing an acidic side chain that likely mimics the electrostatic contribution of the canonical C-terminal carboxylate. Most internal PDZ ligands conformed to modified Class I–like motifs (x[ST]xΦ, where Φ denotes a hydrophobic residue and is called position 0), frequently featuring an xTxFx motif. At the same time, internal ligands belonging to alternative motif consensus classes were also identified, including a WI motif that binds the largely uncharacterized AHNAK2 PDZ domain (**Fig. 3e**), and a WLG motif recognized by ARHGAP23 and ARHGEF12 PDZ domains (**Extended Data Fig. 2**), underscoring the diversity of internal PDZ recognition. Deep mutational scanning (DMS) analysis and alanine scanning of TxF containing SHANK1 PDZ ligands (**Extended Data Fig. 6c**) confirmed the previously reported main motif^45^ and revealed additional contributions of the wild-card and flanking regions. DMS of other selected ligands revealed domain-specific sequence preferences. SYNJ2BP favored a Class I–like motif in which an aspartate at p+1 substitutes for the terminal carboxylate (x[ST]xVDx) (**Fig. 3j**), whereas TAX1BP3 PDZ preferred a more extended motif ([YFW]xx[ST]xΦx; **Fig. 3k**) with reduced dependence on a p+1 aspartate. The upstream aromatic residue (p-5) in TAX1BP3 ligands is likely accommodated by an auxiliary hydrophobic binding site previously implicated in C-terminal ligand binding^46^. Both PARD3B PDZ2 (ExxVD) and PARD6B (Exx[VI][DE][IV]) preferred upstream acidic residues in addition to a p+1 acidic group (**Fig. 3l,m**). Together, these findings reveal internal-motif binding as a widespread feature of PDZ domains, fundamentally expanding the known scope and variety of PDZ-mediated interactions.

### RNA-binding domains recognize SLiMs

While select RNA recognition motif (RRM) domains and other RNA-binding domains have been reported to engage peptide ligands^47^, their SLiM binding has not been systematically explored. To map this landscape, we screened 35 RRM domains for peptide binding, successfully recapturing the canonical UHM-ligand motif (ULM)^48^ recognized by the RRMs of U2AF2, RBM17, and HTATSF1 (**Supplementary Table 2**). For the known peptide-binding RRM domain of the exon-junction complex hub RNPS1^49^ we defined critical binding determinants via DMS of a peptide from its reported interactor HEXIM2^50^ (**Fig. 3n**). We further confirmed binding of 66 other RNPS1-binding peptides by SIMBA, including one from the polyadenylate-binding protein 1 (PABPC1_78-93_) (**Fig. 3o**). We used alanine-scanning peptide arrays to validate motif binding for the uncharacterized peptide-binding RRMs of EIF3B (Wxxx[LF]), RBM7 (SxxLxxAL), and SNRPA (WxxAxxxW) (**Fig. 3p-r**), as well as secondary binding modes for established motif-binding RRMs for ALYREF and PTBP1 (detailed below). AlphaFold3^15^-based modeling (**Fig. 3q-r**) suggests that the peptide-binding pocket in these domains is similar to the previously defined binding surface^48^. Beyond the RRM domains, we found potential ligands of the PCBP1 KH1 domain. Peptide array analysis confirmed robust binding of two overlapping peptides from the RNA binding protein pumilio homolog 1 (PUM1) and of a peptide from the ubiquitin-associated protein 2 (UBAP2; **Fig. 3s**). Together, these findings establish a set of RNA binding domains as dual-function modules that integrate RNA recognition with IDR-mediated protein assembly.

### RRM domains show plasticity of motif binding

The RRM domains from PTBP1 and ALYREF exemplify an additional layer of adaptability: plasticity in recognizing distinct motifs within a common binding site.

ProP-PD derived ligands for the PTBP1 RRM domain showed enrichment of an LLxxP motif (**Extended Data Fig. 2**), which is similar to its previously described LLGxxP consensus (called the PTBP1 RRM interacting motif, PRI^51^) but with a distinct spacing between the leucine residues and the proline. Alignment of selected PTBP1 RRM ligands shows conservation of the glycine at the p3 position and prolines at both p5 and p6 (**Fig. 4a**). Consistent with the previously defined motif, DMS of a model peptide from MATR3 supported a preference for a [GS][ILVM][ILVM]GxxP motif (**Fig. 4b**). Through SIMBA based affinity ranking, we validated binding of 18 ligands (**Supplementary Table 6**), all of which contained the consensus motif except one high scoring peptide from RNF165. DMS of this RNF165 peptide uncovered an alternative double aromatic-containing motif (**Fig. 4c**). We also identified multiple ligands of the mRNA export adaptor ALYREF, including several from proteins involved in mRNA processing and export. Some ligands contained an RLG motif (**Fig. 4d**) similar to previously described viral ligands for ALYREF^52^. This consensus was confirmed by alanine scanning and DMS, which collectively revealed additional N-terminal residue preferences (**Fig 4 e,f,g**). Intriguingly, other ligands contained a distinct [DS][ALV][KR]xx[LM] motif (**Fig. 4h**). This alternative motif class was validated by profiling of additional ligands (**Fig. 4j-k**). The ALKBH5–ALYREF interaction was further validated by co-IP (**Fig. 4l**), which was disrupted by mutation of the DLRxxL motif in ALKBH5. AlphaFold3 modeling placed ligands containing either the RLG or the DLRxxL motifs in a partially overlapping binding site (**Fig. 4m**), and a competition FP-monitored assay supported their mutual exclusivity (**Fig. 4n**). Thus, the ALYREF RRM domain uses a single binding surface to accommodate sequence-divergent motifs, emphasizing the plasticity of peptide recognition.

**Figure. 4.**
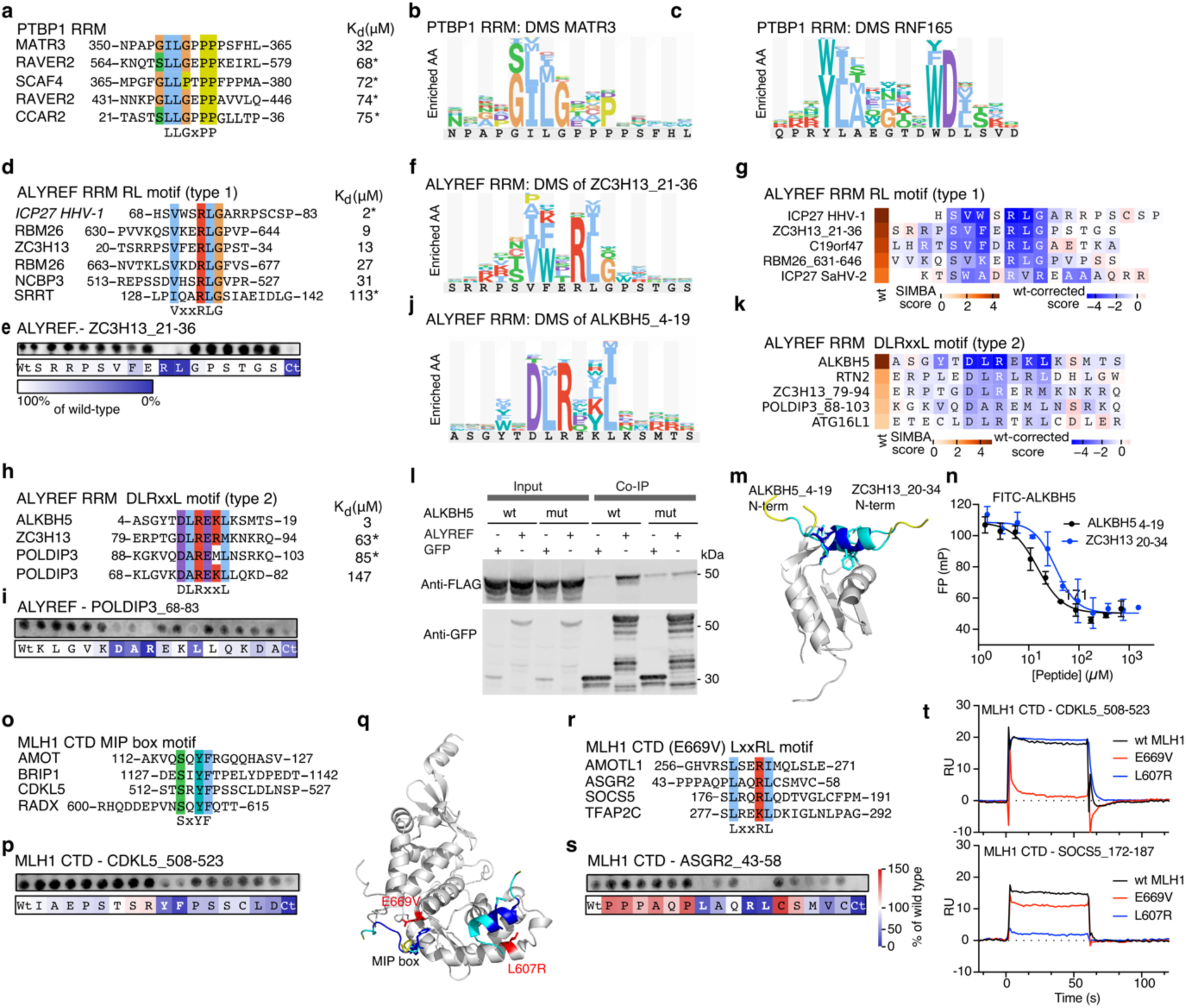
Hidden complexity in SLiM recognition. **a,** PTBP1 RRM ligands with a common LLxxP motif, together with affinities determined by FP assays (n=3) or estimated from SIMBA scores (indicated by *; n=6). **b**, **c,** DMS analysis of two distinct ligands validates the expected motif with some variation (**b**), and discloses an alternative motif in a distinct peptide backbone (**c**). **d–n,** The ALYREF RRM peptide-binding site recognizes two distinct SLiMs. Representative type 1 ϕxxRLG ligands (**d**) and type 2 DLRxxL ligands (**h**) are shown together with affinities determined by competitive FP (n=3) or estimated from SIMBA scores (*; n=6). Alanine-scanning peptide arrays validate the key determinants of the two motifs (**e,i**). PSSMs derived from DMS define the type 1 motif in ZC3H13 (**f**) and the type 2 motif in ALKBH5 (**j**), and SIMBA-based alanine scanning of five type 1 ligands, including two viral ligands (**g**), and five type 2 ligands (**k**) further resolves motif preferences. (**l**) Co-immunoprecipitation of GFP-tagged ALYREF with FLAG-tagged ALKBH5 and a ALKBH5 _11_-REKL-_14_ to _11_-AAAA-_14_ motif-mutant validates the ALYREF RRM type 2 motif-based interactions in cells (n = 3). (**m**) Superimposed AF3 models suggest that the two ligand classes bind the same ALYREF surface in opposite orientations (ALKBH5_4-19:_ ipTM 0.7p, TM = 0.77; ZC3H13_20-34:_ ipTM 0.63, pTM 0.76). (**n**) Peptides of both classes compete with a FITC-labelled ALKBH5 peptide, supporting binding to the same site. **o-t**, MLH1 CTD recognition extends beyond the canonical MIP-box motif through a second cryptic peptide-binding site. Representative ligands recovered with wild-type MLH1 CTD (**o**) or the MIP-box mutant E669V (**r**) define two ligand classes, which are validated by alanine-scanning peptide arrays (**p,s**) using wild-type MLH1 CTD. AF3 models suggest distinct binding modes for MIP-box-(CDKL5_508-523_: ipTM, 0.76, pTM 0.9) and LxxRL-containing (SOCS5_173-188_: ipTM 0.83, pTM = 0.9) peptides (**q**). SPR analysis of immobilized wild-type MLH1 and pocket mutants (E669V and L607R) with representative ligands confirms selective disruption of the two binding sites (**t**). See **Supplementary Figure 4s-v** for additional representations of the modelled complexes.

### Multisite SLiM recognition exemplified by MLH1

The ASHI dataset further reveals versatility in the SLiM-mediated recognition, with numerous domain families harboring multiple distinct binding pockets that expand their interaction capacity. This is illustrated by the identified kinase ligands (**Fig. 2**), and it is further exemplified by the C-terminal domain (CTD) of the DNA mismatch repair protein MLH1. MLH1 recognizes the canonical MIP-box motif (xSx[FY]Fx) via its C-terminal domain^53^. We identified numerous MIP-box-containing peptides, including motifs in known interactors such as BRIP1^54^, CDKL5^2^, and AMOT^55^ (**Fig. 4o**). Affinities were in the low-micromolar range (**Supplemental Table 4**), and motif dependence was confirmed by alanine-scanning peptide arrays (**Fig. 4p**). However, recent work suggested that MLH1 may harbor a second peptide-binding site^56^. We therefore disrupted the canonical MIP-box pocket by a E669V mutation, which was confirmed by FP-monitored affinity measurement (**Supplementary Table 4**). The ProP-PD selection results for the MLH1 CTD E669V mutant were depleted in MIP-box containing peptides (**Supplementary Table 2**) but enriched in other sequences, including an AMOTL1-derived peptide also recovered with wild-type MLH1 (**Fig. 4r**). Alanine-scanning peptide array analysis defined an LxxRL motif (**Fig. 4s**), and AlphaFold3^15^ modeling positioned these peptides in a distinct groove of the MLH1 CTD (**Fig. 4q**). A MLH1 CTD L607R mutation selectively disrupted binding to LxxRL ligands without affecting MIP-box recognition (**Fig. 4t**). Notably, both mutations tested, E669V and L607R, correspond to clinical variants of uncertain significance. Some interactors, such as AMOT, contain both MIP-box and LxxRL motifs. The results support the presence of two discrete peptide-binding sites in MLH1, illustrating a broader principle of multisite recognition among peptide-binding domains.

These examples underscore the inherent versatility of SLiM recognition: domains can accommodate alternative bound conformations within a single pocket or evolve multiple, spatially distinct binding surfaces. This plasticity likely underpins the tunability of interaction networks, allowing weak, transient motifs to encode precise yet flexible regulation.

### IDR context tunes motif specificity

SLiMs do not operate as isolated sequence modules; their affinity and biological specificity are governed by the local sequence landscape of the wider IDR. We further investigated the architectural constraints surrounding human SLiMs to understand how IDRs modulate interaction networks. At the simplest level, most motif-containing proteins contain only one motif instance (61%; 2,531) (**Fig. 5a**). However, individual motifs were not always exclusive to a single binding partner. For example, between 32% and 52% of SH3-, WW- and PDZ-binding motifs were found to bind more than one domain family member from distinct proteins, suggesting a potential for competitive binding (**Fig. 5b).** Nevertheless, most peptides were recovered as ligands for a single domain. This specificity was evident even among domains of the same families with partially overlapping motif preferences, such as the PDZ domains of SYNJ2BP, TAX1BP3, PARD3B and PARD6B. SIMBA-based affinity ranking across these PDZ domains demonstrated selective ligand recognition despite related key sequence features (**Fig. 5c**). DMS of individual ligands of these domains identified gatekeeper residues at several positions that strongly influenced domain discrimination and revealed single substitutions that alter binding profiles (**Fig. 5d**). Thus, subtle sequence variation provides selectivity between closely related domain family members. Moreover, flank-swapping experiments with partially cross-reactive PARD3B and PARD6B ligands showed that residues outside the core motif strongly influence affinity and selectivity **(Fig. 5e-g)**, likely through both direct contacts and modulation of the overall peptide secondary structure propensity. In support of the latter, AlphaFold3 modelling suggests that a favored C-terminal flanking region from the peptide FAM124B promotes β-hairpin formation (**Fig. 5h**), which may facilitate docking into the PDZ binding pocket. Together, these results show that the information encoded in IDRs is layered: SLiM-mediated specificity depends not only on the core motifs, but also on sequence context.

**Figure 5.**
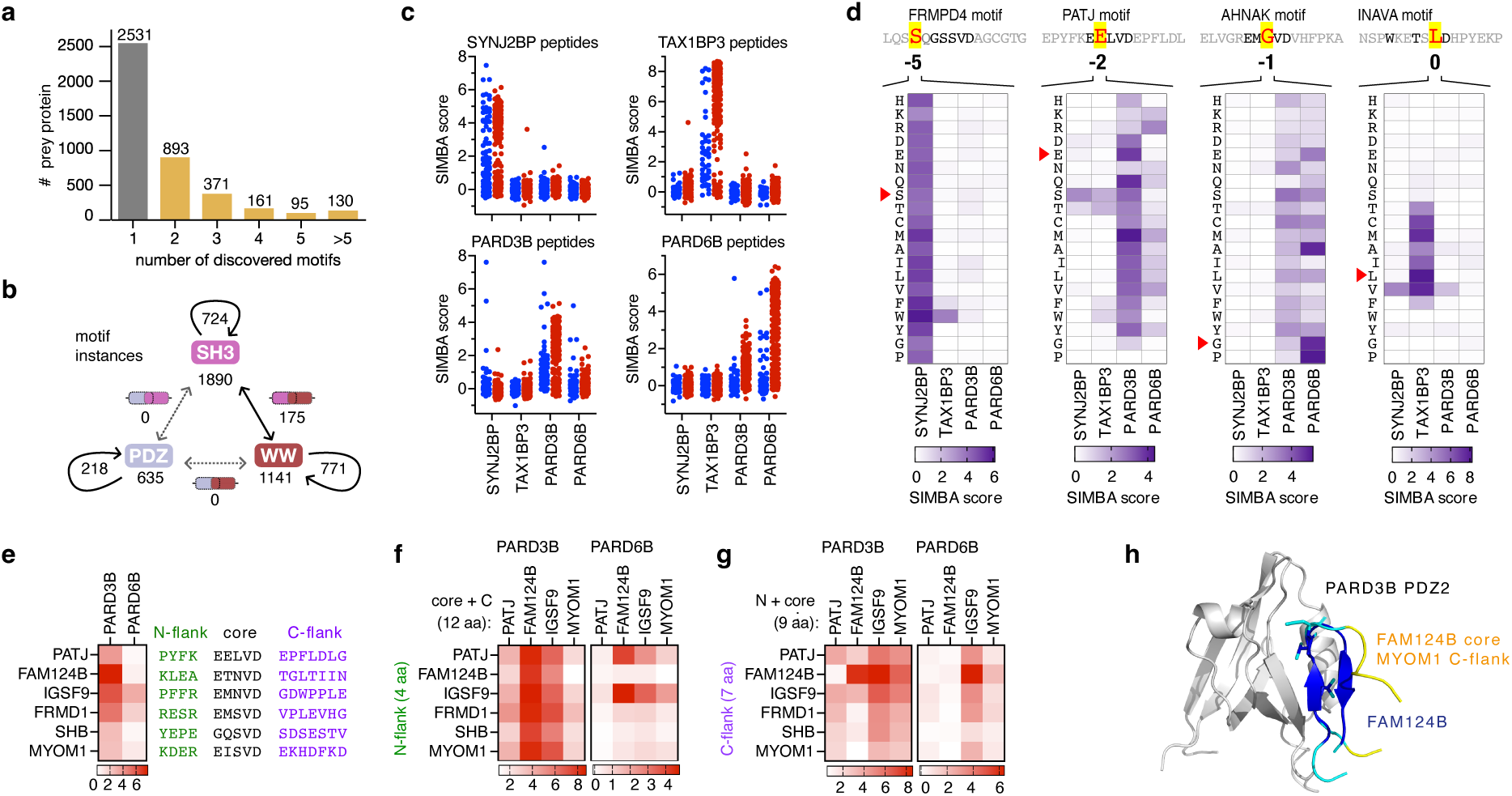
Organizational and sequence features governing specificity versus overlap of SLiM-domain interactions. **a,** Number of motifs found per prey protein. **b**, Competition among PDZ-, SH3- and WW-binding motifs within domain families (curved arrows) and between domain families (straight arrows). Numbers indicate the total motif instances detected for each family, the subset bound by multiple members of the same family, and regions containing overlapping binding sites for different domain families. **c,d**, Specificity despite overlapping motif preferences. **c**, SIMBA-based all-by-all affinity ranking of ligands across the PDZ domains of SYNJ2BP, TAX1BP3, PARD3B PDZ2 and PARD6B. Both ProP-PD-derived ligands (blue) and ligands generated from SIMBA based DMS of selected parental peptides are shown (see **Supplementary Table 5**). **d,** SIMBA-DMS analysis revealing position- and domain-specific gatekeeper residues that shape ligand selectivity across the four PDZ domains (n = 6); red arrowheads denote wt residues. **e–h,** Context-dependent specificity**. e,** Sequences and SIMBA scores of six parental PARD3B PDZ2 ligands used for flank-swapping experiments with PARD3B PDZ2 and PARD6B (n = 6). **f**, Exchange of the N-terminal flank selectively altered affinity for PARD6B, as illustrated by the high-affinity combinations generated by pairing the N-terminal flanks of PATJ and IGSF9 with the motif core and C-terminal flank of FAM124B. **g**, Exchange of the C-terminal flank affected affinity for both PARD3B PDZ2 and PARD6B. **h**, AlphaFold3 modelling of PARD3B PDZ2 and two variants of the FAM124B peptide (wt (ipTM 0.89, pTM 0.88) and FAM124B with the C-terminal region of the MYOM1 peptide (ipTM 0.69, pTM 0.74) suggests that the context influences the secondary-structure propensity of the bound peptide; the FAM124B C-terminus may promote a beta-hairpin that depends on a Gly residue at p+3 (**Supplementary Figure 5c**). **See Extended Data** Figure 4w**,x** for additional representations of modelled complexes.

This conclusion was reinforced by testing whether core motif matches alone could predict binding. Among 222 sequences predicted as candidate ligands based on matches to the consensus motif for one of nine bait domains, only 44 (20%) scored as binders when tested by SIMBA (**Supplementary Table 6**). These results demonstrate that the broader sequence context contributes to distinguishing binders from non-binders.

### Specificity and interplay through competition and motif co-occurrence

The ASHI dataset allowed us to examine how multiple motifs are combined within the IDRs of individual proteins. Notably, 39.5% of identified prey proteins contained two or more motifs (**Fig. 5a**). We identified three distinct modes of multi-motif organization. The first mode involves overlapping motifs that bind different domain families (Fig. **5b**, **6a**). This set of 547 overlapping motifs is dominated by polyproline-rich peptides that are bound by SH3, WW and EVH1 domains **(Supplementary Table 2**), suggesting that these regions may function as low-specificity competitive hubs.

**Figure 6.**
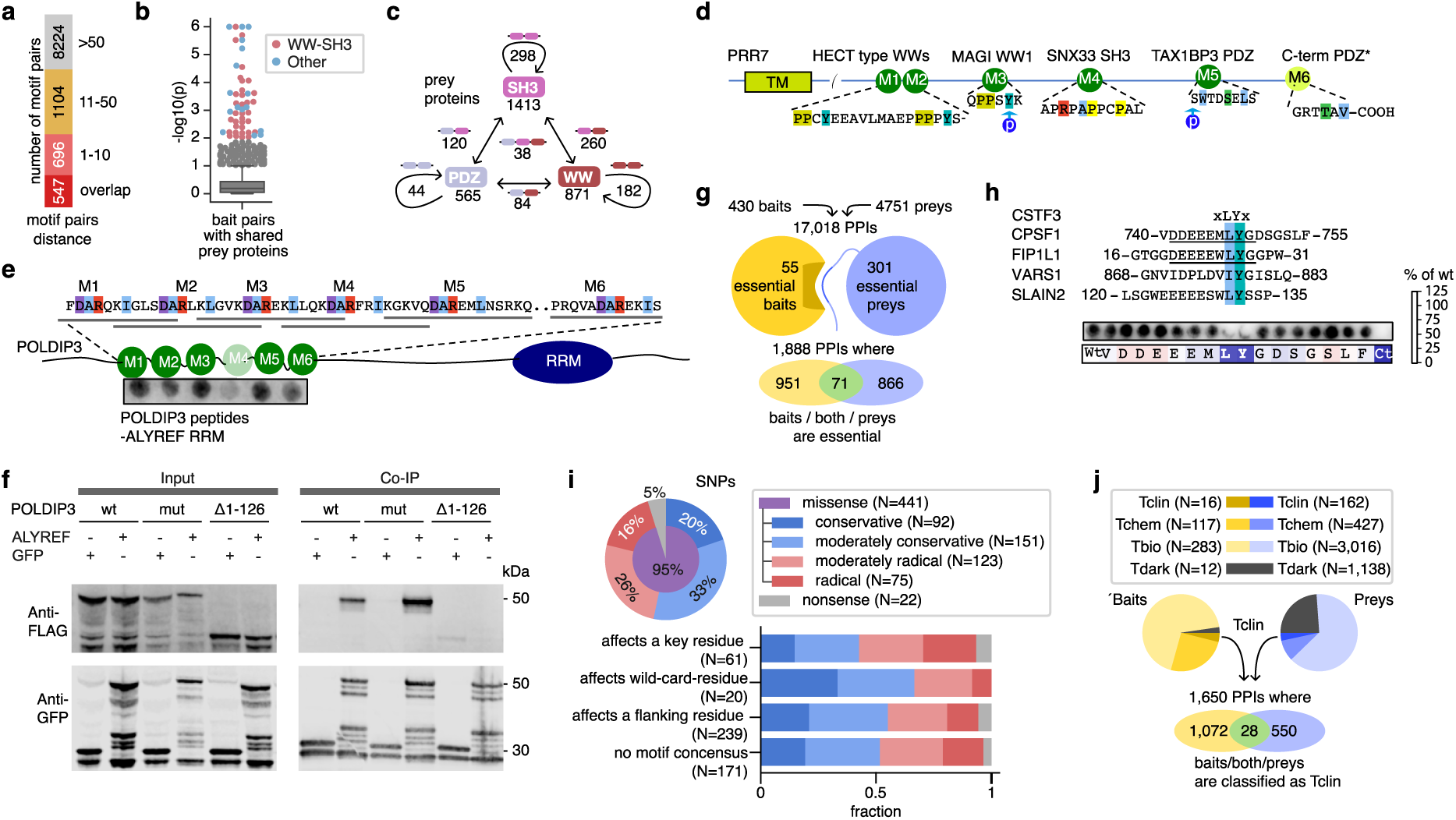
Motif co-occurrence and arrays reveal principles of SLiM-mediated recognition. **a**, Overview of the number of amino acids to the nearest motif neighbor in prey proteins with more than one type of distinct motif. **b,** Prey protein co-occurrence analysis showing that pairs of SH3 and WW domains from specific bait proteins frequently bind to the same prey proteins (**Supplementary Table 8**). **c**, Overview of prey proteins containing non-overlapping PDZ-, SH3- and WW-binding motifs. Curved arrows denote co-occurrence within the same domain family, straight arrows denote co-occurrence between different domain families, and the central value denotes the 38 prey proteins containing motifs recognized by all three families. **d–f**, Motif arrays in SLiM-mediated recognition. **d,** WW-binding [LP]PxY motifs frequently co-occur at close distance, as exemplified by the transmembrane protein PRR7, which also contains SH3 and PDZ binding motifs. **e**, The N-terminus of POLDIP3 contains an array of six identified or predicted ALYREF RRM-binding motifs, five of which were validated by peptide-array analysis. **f,** Co-immunoprecipitation of FLAG-tagged ALYREF with full-length wild-type GFP-tagged POLDIP3 and a motif-mutant of the third motif (M3 in panel **e**) or truncated variants, supporting the importance of the motif-array for binding in the context of the full-length protein (n = 3). **g,** Overview of the number of PPIs in the ASHI dataset that involve essential bait or prey proteins. **h,** Alignment of representative peptides binding to the essential protein CSTF3 (top) together with validation of a xLYx consensus motif through peptide SPOT array analysis. The binding peptides overlap with essential regions in CPSF1 and FIP1L1 (underscored)^63^. **i,** SNPs distributions in ProP-PD enriched ligands (top) and at motif feature level (bottom), classified as nonsense (STOP introducing mutation) and missense mutations, with missense changes subclassified on base of the change Grantham score. **j,** Distribution of Pharos^68^ classification of ASHI bait and prey proteins, together with the number of PPIs involving Tclin classified baits and/or proteins. For Tclin proteins, at least one approved drug exists.

A second mode involves co-occurrence of non-overlapping motifs that bind different domain families. Pairs of WW- and SH3- domains frequently bound to IDRs of the same prey proteins (**Fig. 6b**). In some cases, these motifs occur together with PDZ-binding motifs (**Fig. 6c**), as exemplified by the proline-rich protein 7 PRR7 (**Fig. 6d**), a membrane-anchored protein that is largely unstructured. Thus, members of the three major peptide-binding domain families can converge binding on the same prey proteins by binding motifs positioned at different sites within a single IDR.

A third mode involves IDRs containing repeated motifs recognized by the same domain family. This architecture is well suited to engage proteins that form homo-oligomers or that contain multiple copies of the peptide-binding domain, which can increase effective affinity and selectivity through multivalency and avidity. Alternatively, repeated motifs can increase the apparent affinity for a single domain by providing an increased local concentration. These architectures are particularly evident for prey proteins interacting with WW domains proteins, representing 28% of the prey proteins with 2 or more homotypic motifs (**Supplementary Table 2**). PRR7 again illustrates this principle, as it harbors tandem WW domain-binding motifs found to engage the WW3 and WW4 domains of WWP1, as well as WW domains from other HECT-type E3 ligases (**Fig. 6d**). The short linker between these tandem motifs is compatible with multivalent binding, based of AF3 modelling of the complex (**Extended Data Fig. 5y**), and is in line with the existing literature on multivalent binding of WW motifs^57–60^. Thus, the spacing of the linked motifs, and their respective binding domains, emerges as a likely important contributor to the binding strength and specificity of their interactions.

Arrays of motifs were also found for other domain types, such as the ALYREF RRM domain, for which the ProP-PD screening identified three binding motifs in the N-terminal region of polymerase delta-interacting protein 3 (POLDIP3), and two motifs each in the proteins RBM26 and ZC3H13 (**Fig 4d-n**). Further sequence analysis revealed that the POLDIP3 N-terminal region is particularly dense with ALYREF binding motifs, containing six consensus motifs of which five showed detectable binding (**Fig. 6e**). Co-immunoprecipitation showed robust interaction of the full-length proteins (**Fig. 5n**) although the individual POLDIP3-derived peptides bound weakly (80-150 μM; **Supplementary Tables 4, 6**). The interaction was not affected by mutating a single binding site (**Fig. 6f**), but it was abolished by truncating the POLDIP3 N-terminus, consistent with multiple low-affinity sites collectively stabilizing the interaction. In a cellular context, this could be accomplished by multimerized ALYREF in the TREX complex^61^ serving as a multivalent scaffold for POLDIP3 binding. Thus, the organization of SLiM information in the IDRs ranges from the presence of relatively simple interaction elements to more complex motif arrays.

Together, these findings show that SLiM information in IDRs is layered at multiple scales. At the level of individual motifs, specificity is tuned by residues within and around the core binding site. At the level of the full IDR, interaction output is shaped by motif overlap, co-occurrence, spacing and repetition. Thus, IDRs encode interaction specificity not only through individual SLiMs, but through the organization of motif ensembles that can promote competition, avidity and multi-partner recruitment.

### The ASHI dataset links core cellular functions to disease and therapeutic opportunities

The here generated ASHI data set comprises more than 20,000 motif–domain interactions (**Supplementary Table 2;** here generated plus previously reported), corresponding to 17,018 putative protein–protein interactions. The interacting proteins span diverse functional classes, including membrane signaling, cytoskeletal regulation, RNA metabolism, DNA repair and protein quality control, highlighting the broad cellular reach of SLiM-based recognition and their links to core cellular processes. Notably, over 10% of the protein-protein interactions involve proteins where either the bait or the prey is classified as essential in DepMap^62^ (**Fig. 6g**). A subset (71) of interactions connects two essential proteins, such as the binding of the ALYREF RRM domain to a SLiM in SRRT (**Fig. 4d**). In addition, 62 of the SLiM-containing IDRs overlap with regions previously implicated in cell proliferation^63^. These include essential motifs in CPSF1 and FIP1L1 that bind to the protein CSTF3, for which our platform independently mapped the precise molecular determinants (**Fig. 6h**). ASHI also identified the binding partners for uncharacterized yet essential motifs, such as the nuclear pore complex-interacting protein family member B4 (NPIPB4) binding the nuclear export pathway protein RANBP2 PH domain (K_d_ = 1.7 µM, **Extended Data Fig. 3**).

Beyond fitness, ASHI systematically rationalizes disease-associated variation in the disordered regions. We identified 436 disease-associated variants in SLiM-containing peptides from 215 proteins (ClinVar^64^, COSMIC^65^). Only 5% of the single nucleotide polymorphisms (SNPs) with a reported pathological effect introduce a STOP codon. The remaining SNPs were distributed broadly across both key motif residues and flanking sequences (**Fig. 6i**). Mutations affecting motif core residues had the highest fraction of radical changes, which are likely to disrupt binding. The N-terminal cytoplasmic region of the organic cation/carnitine transporter 2 (OCTN2; _5-_DEVTAFLGEWGPFQR_-19_) contains three motifs recognized by distinct domain classes (the SNX27 PDZ domain, the GGA3 EAR domain, and the cyclophilin-type prolyl isomerases PPIA and PPIE). Pathogenic variants clustering in this single region result in renal carnitine transport defects^66^. These observations support the view that disease-associated variation can perturb SLiM function through disruption of motif cores or flanking regions^10^. Finally, cross-referencing ASHI with drug-target resources further underscored its biomedical relevance: 16 bait proteins (**Fig. 6j**) and 162 prey proteins are established drug targets (**Supplementary Table 2**). While targeted baits are enriched in kinases, clinically annotated prey proteins are enriched in ion channels. Together, these findings position ASHI as a resource for linking IDRs to cellular fitness, disease mechanisms and therapeutic opportunity.

## Discussion

We identified over 20,000 motif-domain interactions by screening more than 800 human protein domains against a million-peptide library spanning the IDRs of the human proteome. This substantially expanded the known human SLiM interactome, providing the most extensive experimental map of SLiM-mediated recognition across the human proteome to date. ASHI shifts the field from a collection of motif family case studies to a proteome-scale view of a major, and previously underexplored, layer of the interactome. Large-scale interaction mapping efforts have historically been optimized for different interaction classes: HuRI^2^, based on yeast two-hybrid, captures direct binary interactions, while BioPlex^4^, based on AP-MS, excels at defining stable complex components. ASHI complements these resources by capturing the weak, transient, motif-mediated interactions that are often missed by traditional approaches. While most large-scale interaction maps are biased towards high-affinity interfaces and stable complexes, ASHI provides the lower-affinity interactions characteristic of SLiMs. This adds a critical, context-dependent organizational layer to the human interactome, superimposing dynamic events onto the more rigid architecture of macromolecular assemblies.

A central conclusion of this study is that SLiM-mediated recognition is more widespread across protein domain space than previously appreciated. Beyond expanding the repertoires of classical peptide-binding modules, ASHI reveals a broader landscape of SLiM-binding across enzymes, chaperones and RNA-binding domains, and identifies unmodified SLiM ligands for domains canonically associated with PTM-dependent binding, suggesting that moonlighting SLiM-binding ability is not an anomaly, but a widespread and underappreciated property across diverse domain folds. The internal motif recognition by PDZ domains also underscores this point. Collectively, these observations suggest that the capacity for peptide recognition evolves repeatedly across diverse domain folds, and that the current catalogue of peptide-binding domains represents only a fraction of those present in the proteome. This, in turn, makes a case for the systematic screening of all structurally autonomous domains in the human proteome for peptide-binding capacity, an achievable goal given current proteome-scale technologies.

ASHI further supports an expanded set of molecular rules of SLiM-based recognition. Core motifs are necessary for binding but are often insufficient to explain specificity. Instead, specificity emerges from the interplay between the motifs and their broader sequence context, including wildcard positions, flanking residues, alternative binding modes and, in some cases, ligand conformational propensity. At the same time, the effective affinity of many interactions will be increased by multivalency, through repeated motifs and multisite recognition, indicating that IDRs often function not as simple carriers of individual motifs, but as densely encoded interaction platforms. These findings substantiate and expand on the current understanding^1,67^. In this view, SLiMs are best understood not as isolated functional modules but as contextual binding sites whose specificity, selectivity and affinity are optimized by sequence context, plasticity and combinatorial organization. These findings help explain how short, degenerate motifs can nevertheless encode selective interactions. In addition, many motif containing proteins are also motif binding, which adds additional layers of complexity.

These macro-level principles could only be resolved through proteome-wide screening across a large and diverse cohort of domains and domain classes. Small-scale studies have been essential for defining individual motifs and domain–ligand mechanisms, but only the large-scale approach used here can reveal the breadth of SLiM recognition, the prevalence of moonlighting peptide-binding activities, and the importance of plasticity, multivalency and multispecificity across the proteome. Together, these findings expose general principles of how IDRs encode interaction information. The scale and diversity of ASHI also positions it as an unparalleled training and benchmarking resource for machine learning models aimed at predicting SLiM-mediated interactions from sequence, which have historically been starved of systematically collected negative and positive experimental data. Furthermore, the representation of essential proteins, disease-associated variants and drug-target-linked proteins within this map directly links this transient interaction layer to cellular pathology and therapeutic opportunity. While ASHI reveals a vast interaction landscape, our study has some inherent limitations. By utilizing isolated domains and short peptides, the screen may not fully capture the effects of full-length protein architecture, PTMs, localization, competition or higher-order assembly. Consequently, some identified interactions may be conditional, while those requiring broader cellular context may remain undetected. Furthermore, because our data analysis is conservative, the 20,000 interactions reported here likely represent a lower bound on the true scope of SLiM-mediated recognition across the screened baits, a conclusion supported by the high validation rates of 46% by the orthogonal SIMBA approach and 93% by fluorescence polarization. While *in vitro* biophysical binding does not always equate to *in vivo* physiology, this dataset provides a tractable roadmap for systematic functional validation. This challenge is substantial given the scale of the dataset but is increasingly tractable as emerging high-throughput functional assays provide new routes to systematic motif characterization. Ultimately, ASHI establishes SLiM-mediated recognition as a widespread organizing principle of the human interactome.

## Data availability

The NMR spectra (FIDs) are available, via the open data repository Zenodo (DOI:10.5281/zenodo.17917033). The HD2 library design is also available via Zenodo (DOI:10.5281/zenodo.21071793). ProP-PD results are available through the ProP-PD portal (https://slim-tools.org/proppd/proppd_data_viewer?library=HD2). High confidence SLiM-domain interactions have been deposited in the MoMaP database (https://slim-tools.org/momap/article?pmid=madhu_2026). PPI data will be deposited through IntAct.

## Acknowledgements

We are grateful for constructs made available by Addgene as listed in Supplemental Table 1, and project students that have contributed with technical support. We thank Jakob Nilsson for providing BUB1, Haribabu Arthanari and Thibault Viennet for providing MED15, Edward A FitzGerald for providing KDM1, Nadine Myers for support with SMYD2 and SMYD3, Hong Zeng and Serah W. Kimani for protein purification of WD40 domains, and Eldar Abdurakhmanov and Helena Danielson for advice and access to SPR Biacore 3000.

## Funding

This work was supported by grants from the Swedish Research Council (Y.I.: 2020-03380, 2024-04306; M.E.: 2024-05496, 2022-06628, 2020-03431). A.K., M.V., H.K., N.M. and J.K.V. were supported by funding from the European Union’s Horizon 2020 research and innovation programme under the Marie Skłodowska-Curie grant agreement No. 860517 (UBIMOTIF), S.L. and C.K. were supported by grant agreement No. 675341 (PDZnet), and A.Z. was supported by No 101119633 (IDPro).

Work by M.S.S, M.J.W, and P.M.P. was funded by grants from the NIH to P.M.P. (R01GM057769 and R01GM145795). N.E.D. was supported by a Cancer Research UK Senior Cancer Research Fellowship (C68484/A28159) and the American Lebanese Syrian Associated Charities.

Sequencing for ProP-PD experiments were performed by the SNP&SEQ Technology Platform in Stockholm. The facility is part of the National Genomic Infrastructure (NGI) Sweden and Science for Life Laboratory and is also supported by the Swedish Research Council and the Knut and Alice Wallenberg. This project made use of the NMR Uppsala infrastructure, which is funded by the Department of Chemistry for Life Sciences and the Disciplinary Domain of Medicine and Pharmacy, Uppsala University and SciLifeLab, and is part of the national Swedish research infrastructure SwedNMR, funded by the Swedish Research Council through grant agreement no. 2021–00167.

SK and AK are grateful for support of the German Cancer Aid (DKH, TACTIC-70115201) and the German Cancer Consortium (DKTK) at the German Cancer Research Center (DKFZ). SK would also like to acknowledge support of the DFG (German Research Foundation) under Germanýs Excellence Strategy – EXC 2026, Cardio-Pulmonary Institute, Project ID: 390649896. CA, SK, SS, AK, LH would like to acknowledge funding by the Structural Genomics Consortium (SGC) a registered charity (no: 1097737) that receives direct member funding from Amgen Inc., Janssen Pharmaceutica NV, and Bristol-Myers Squibb Company, as well as grant funding from the Innovative Health Initiative Joint Undertaking (IHI JU; LIGAND-AI, grant agreement No. 101252959), and the Gates Foundation. Through the IHI LIGAND-AI grant, SGC also receives financial and in-kind contributions from Pfizer Inc., AstraZeneca UK Limited, and Novo Nordisk A/S, Abcam Limited, Chemspace LLC, Enamine Germany GmbH, IBM Research Israel – Science and Technology Ltd., Nuvisan ICB GmbH, Thermo Fisher Scientific (Bremen) GmbH, The Hospital for Sick Children, and Vernalis (R&D) Limited.

## Author contributions

P. M., C.B., L.S., J.K., L. G. L., M. V.: Investigation, Visualization, Data analysis.

S. L.-L., S. L.: Investigation, Data analysis.

I. K.: Visualization, Data analysis.

H. M. K., L. E., Data analysis.

M.S.S., M.J.W., D.S., W. T. P. D, A. Z., A. K., F. M., J. K. V., L. P., A. K., K. G., G. L. R., C. K., R. X.: Investigation,

O. S.-F., S. K, L. H., V. S., C. A, M. E., A. S, R. V.: Supervision, Funding acquisition.

P. M. P.: Investigation, Visualization, Supervision, Funding acquisition. Writing – original draft.

N. E. D., Y. I.: Conceptualization, Investigation, Visualization, Supervision, Funding acquisition. Writing – original draft.

## Competing interests

N. E. D. is a scientific co-founder of RIME therapeutics. The rest of the authors declare that they have no competing interests.

## Supplementary Tables

Supplementary Table 1. Overview of

Supplementary Table 2. ProP-PD data scored.

Supplementary Table 3. Motif statistics.

Supplementary Table 4. FP affinity data

Supplementary Table 5. SIMBA all results

Supplementary Table 6. SIMBA data affinity ranking

Supplementary Table 7. NMR assignments

Supplementary Table 8. Motif-co-occurrence analysis.

Supplemental Table 9 SIMBA_plasmids_oligos

## Extended Data for

**Extended Data 1.**
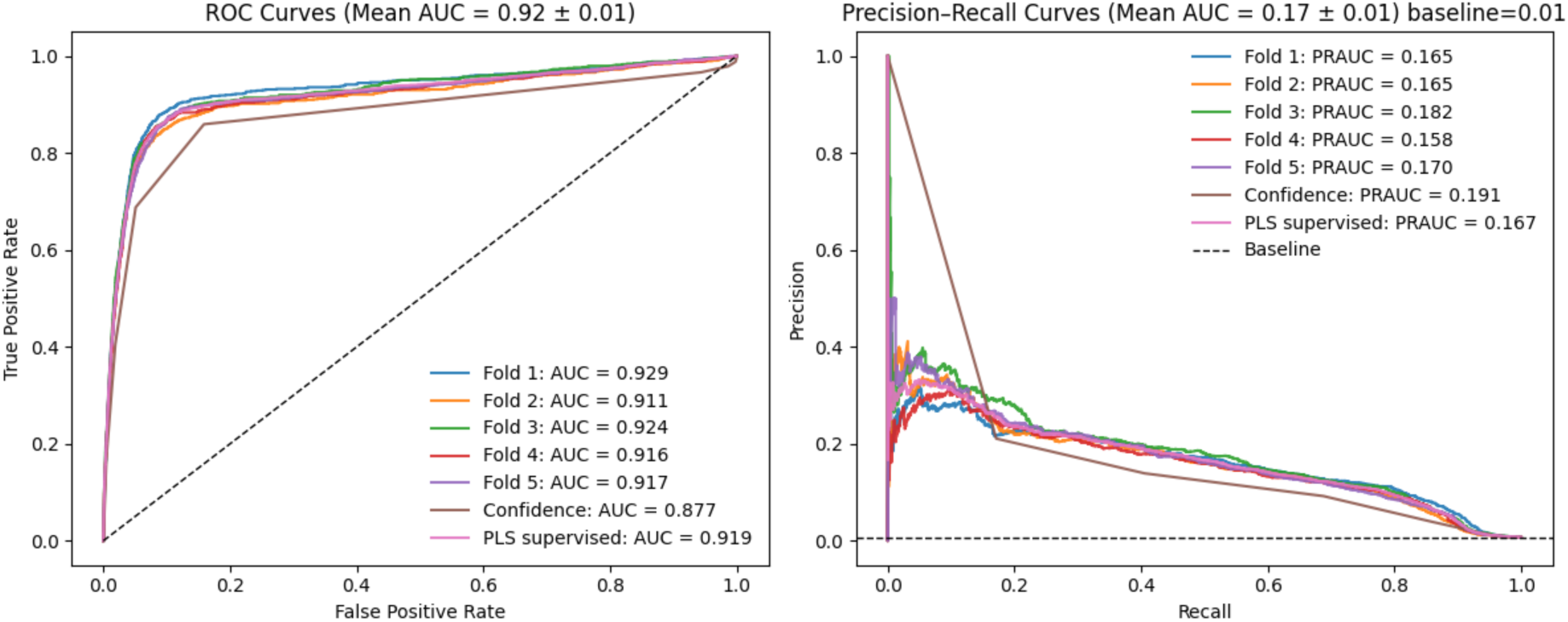
Benchmarking of the partial least squares (PLS) supervised scoring. ROC and Precision-Recall curves comparing five-fold cross-validation of PLS supervised scoring, a model trained on the full dataset, and the legacy confidence score as a baseline comparator. For details see Supplementary Methods.

**Extended Data Fig. 2.**
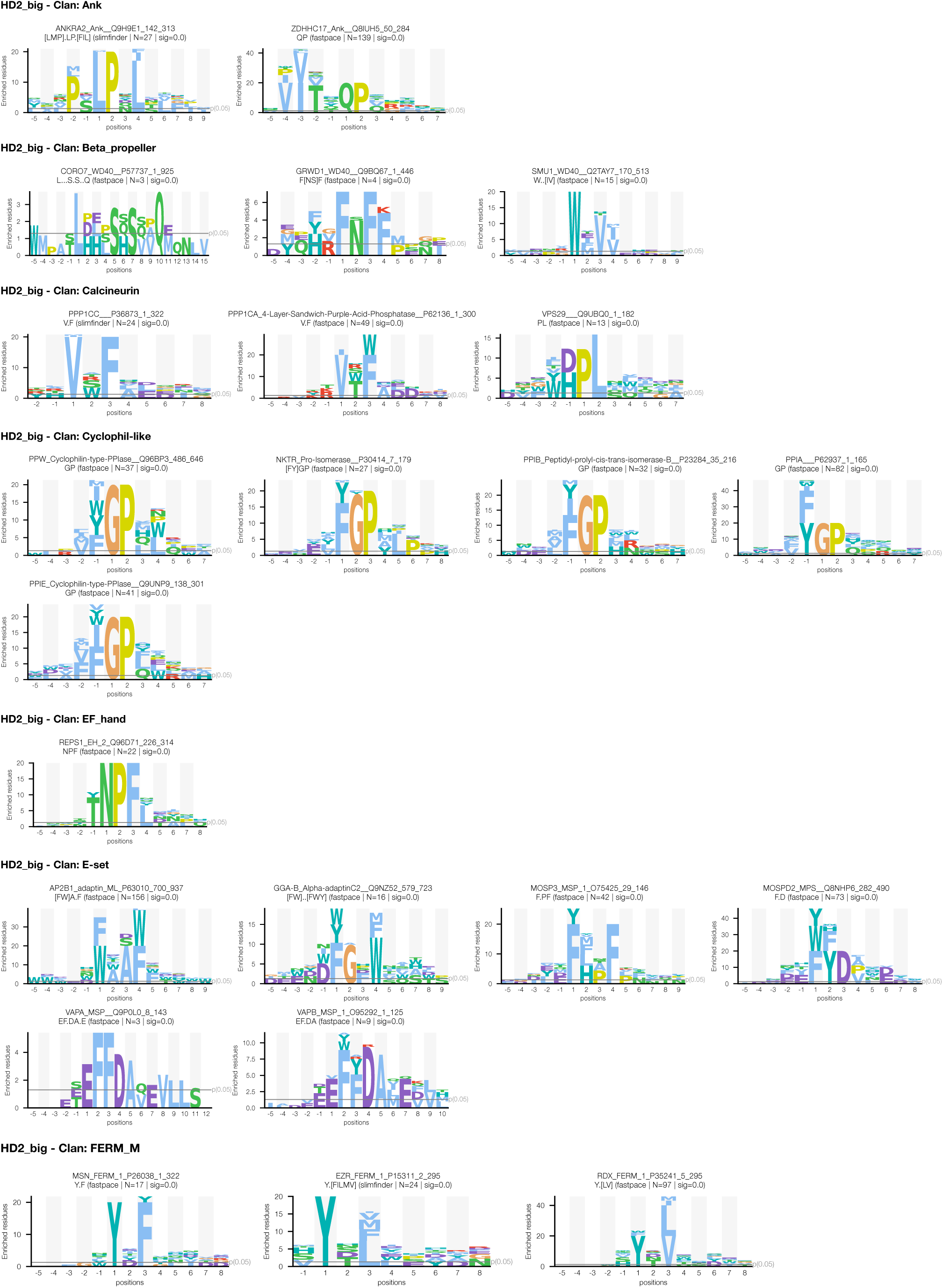
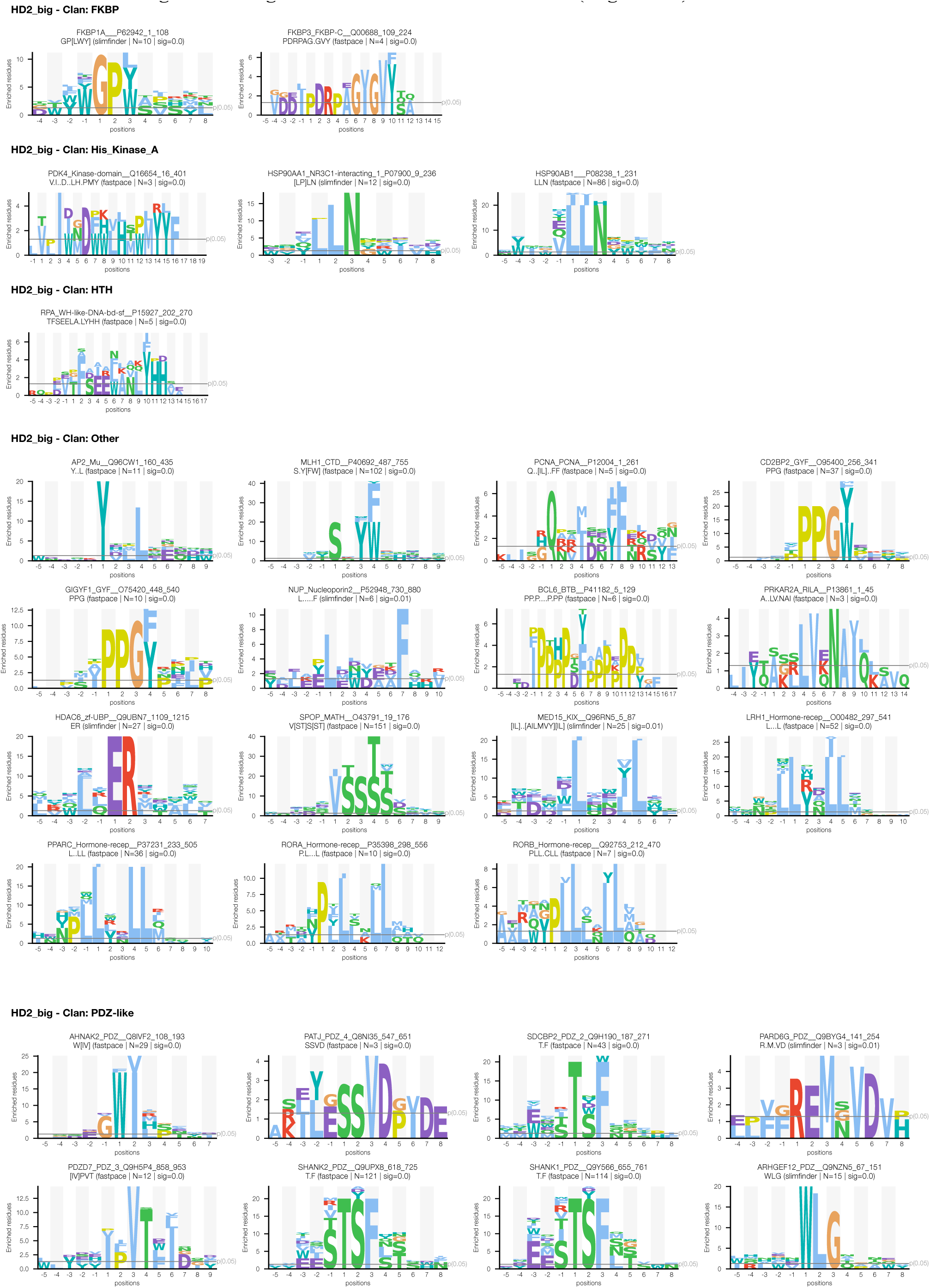
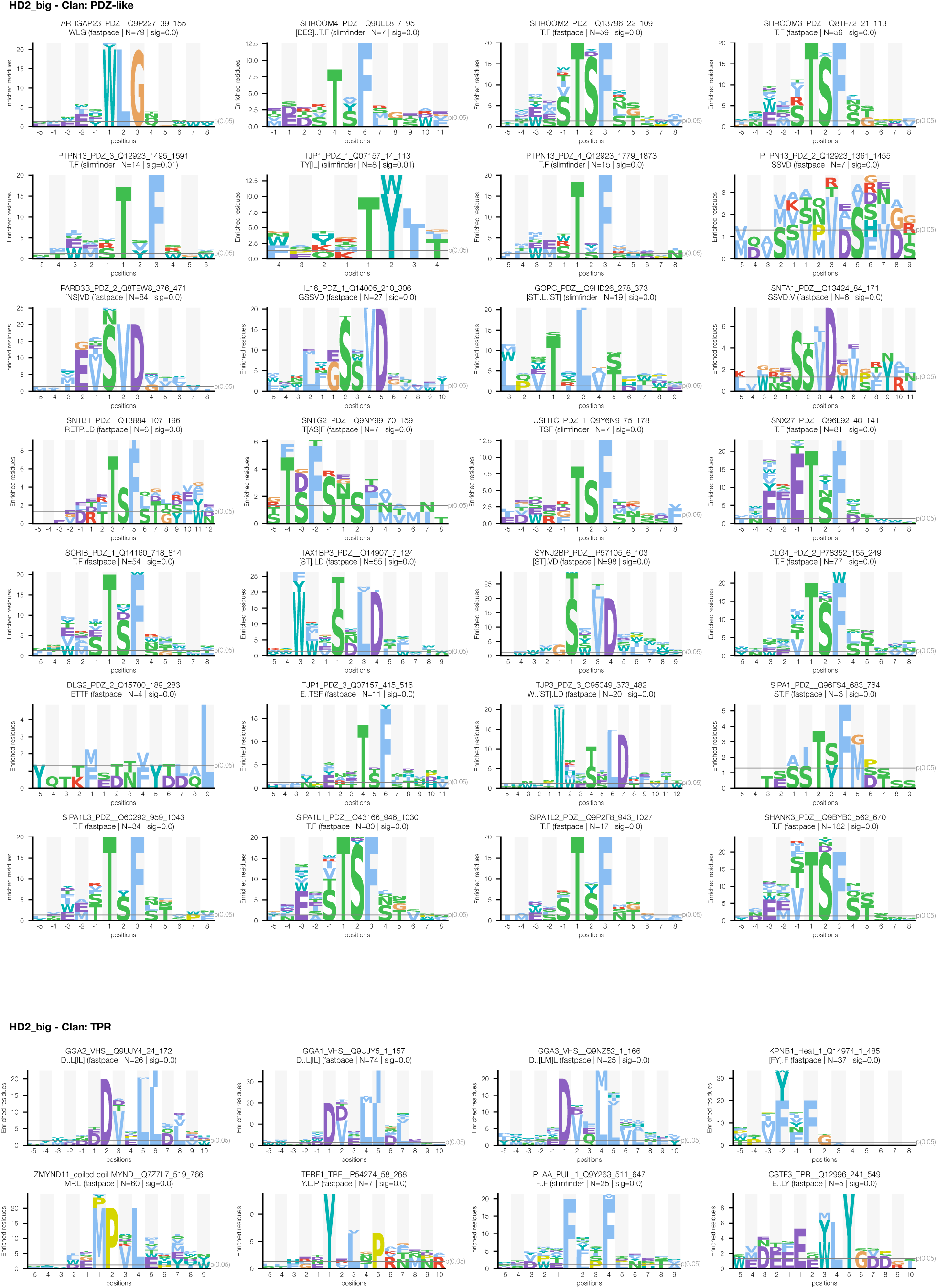
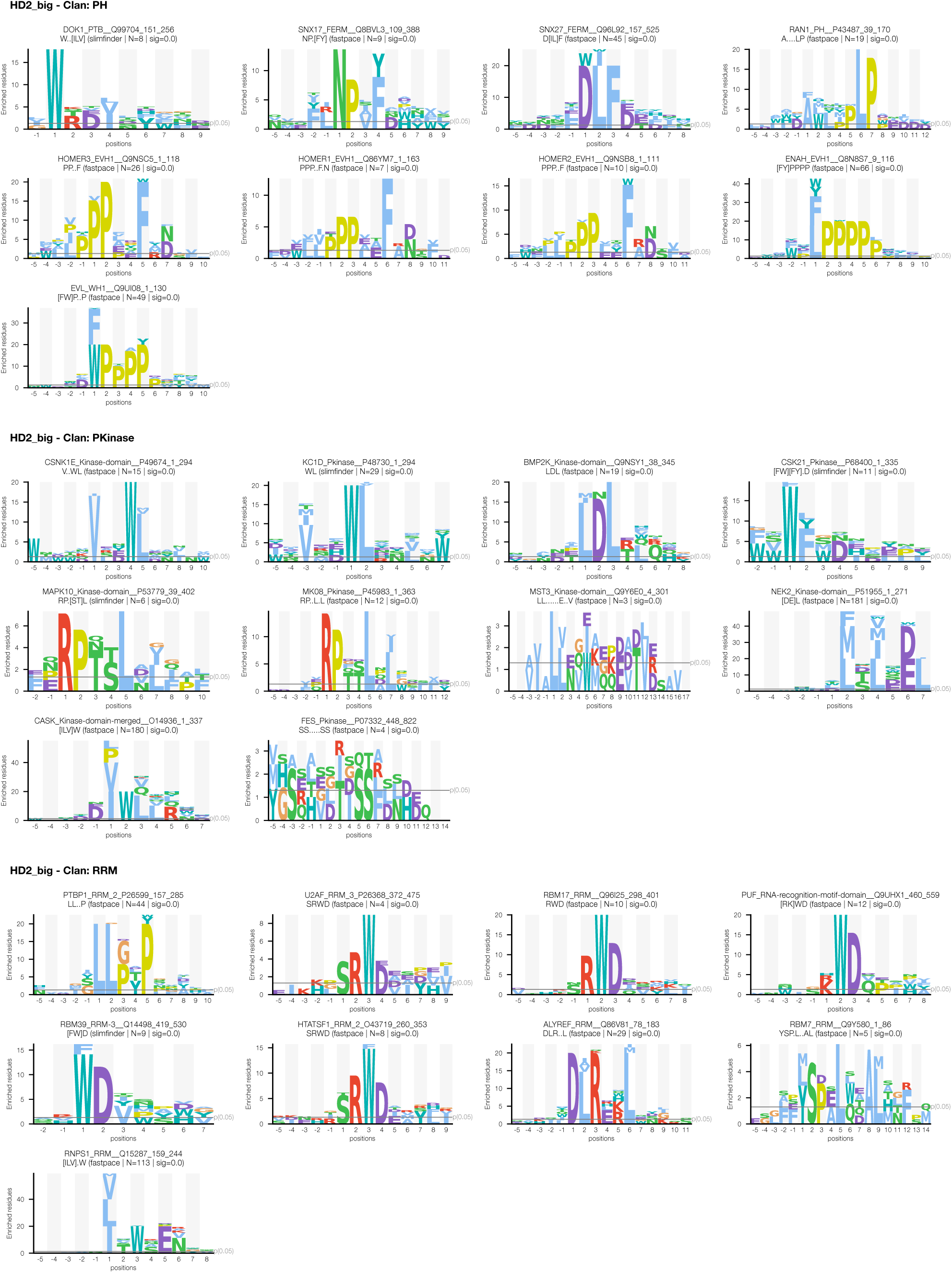
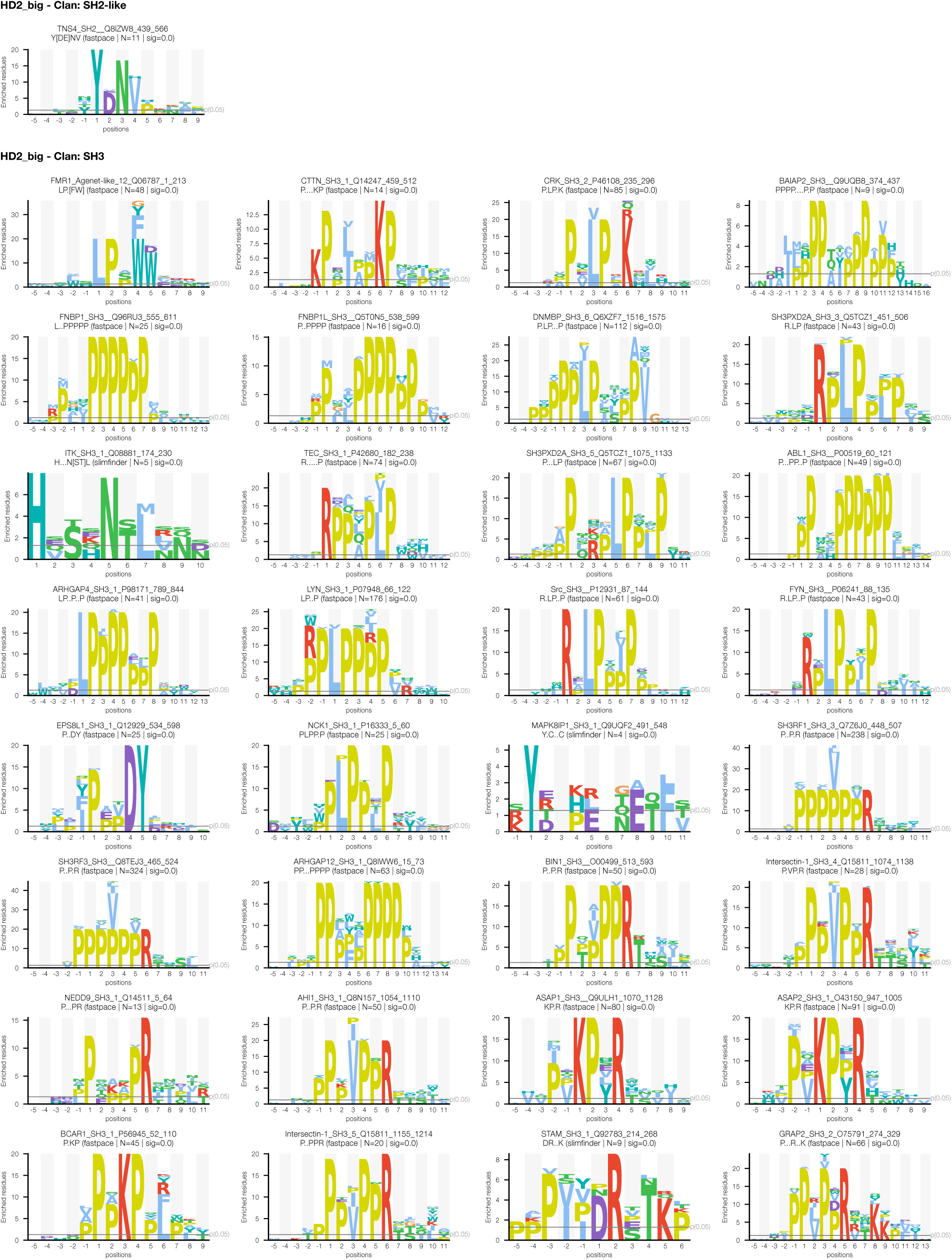
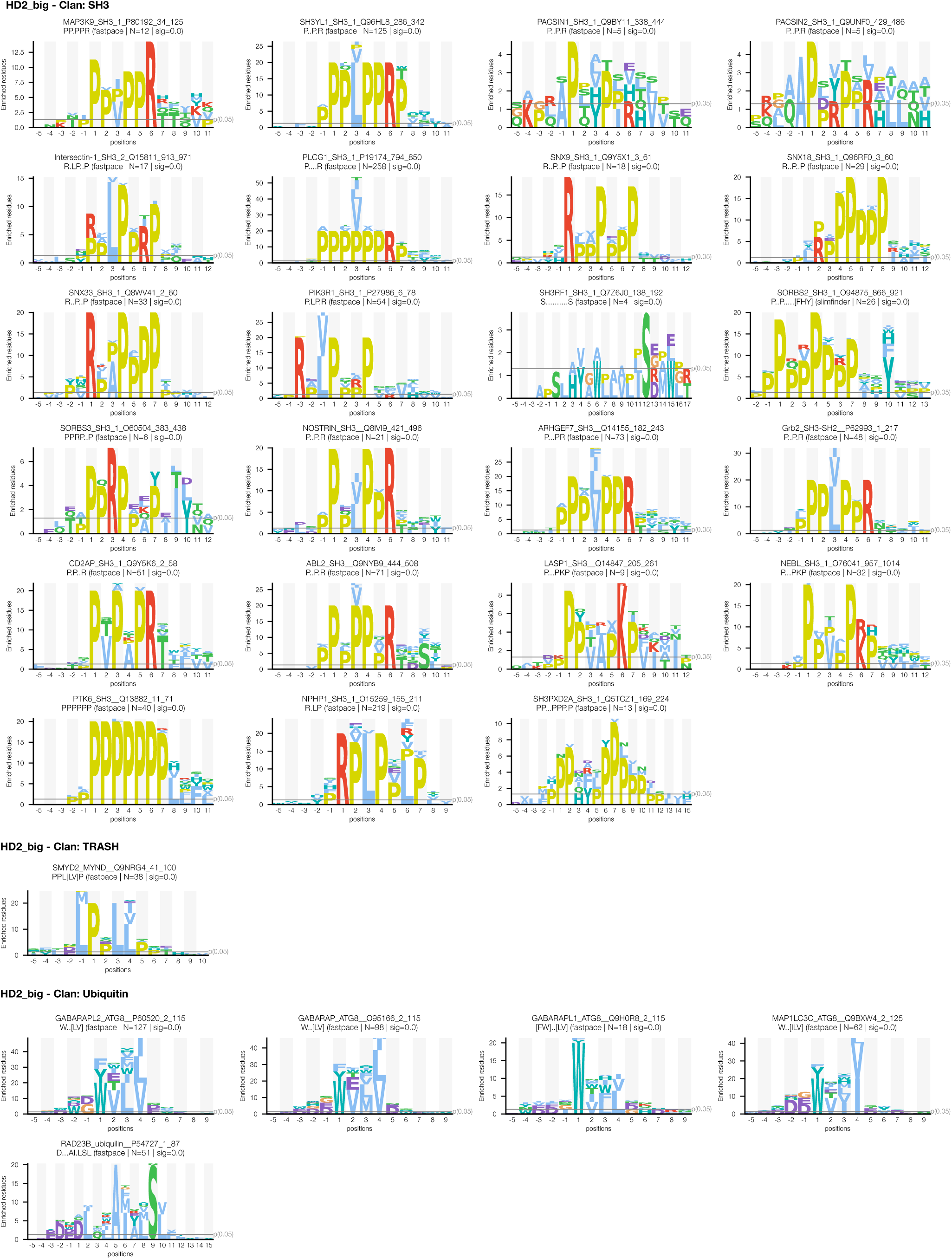
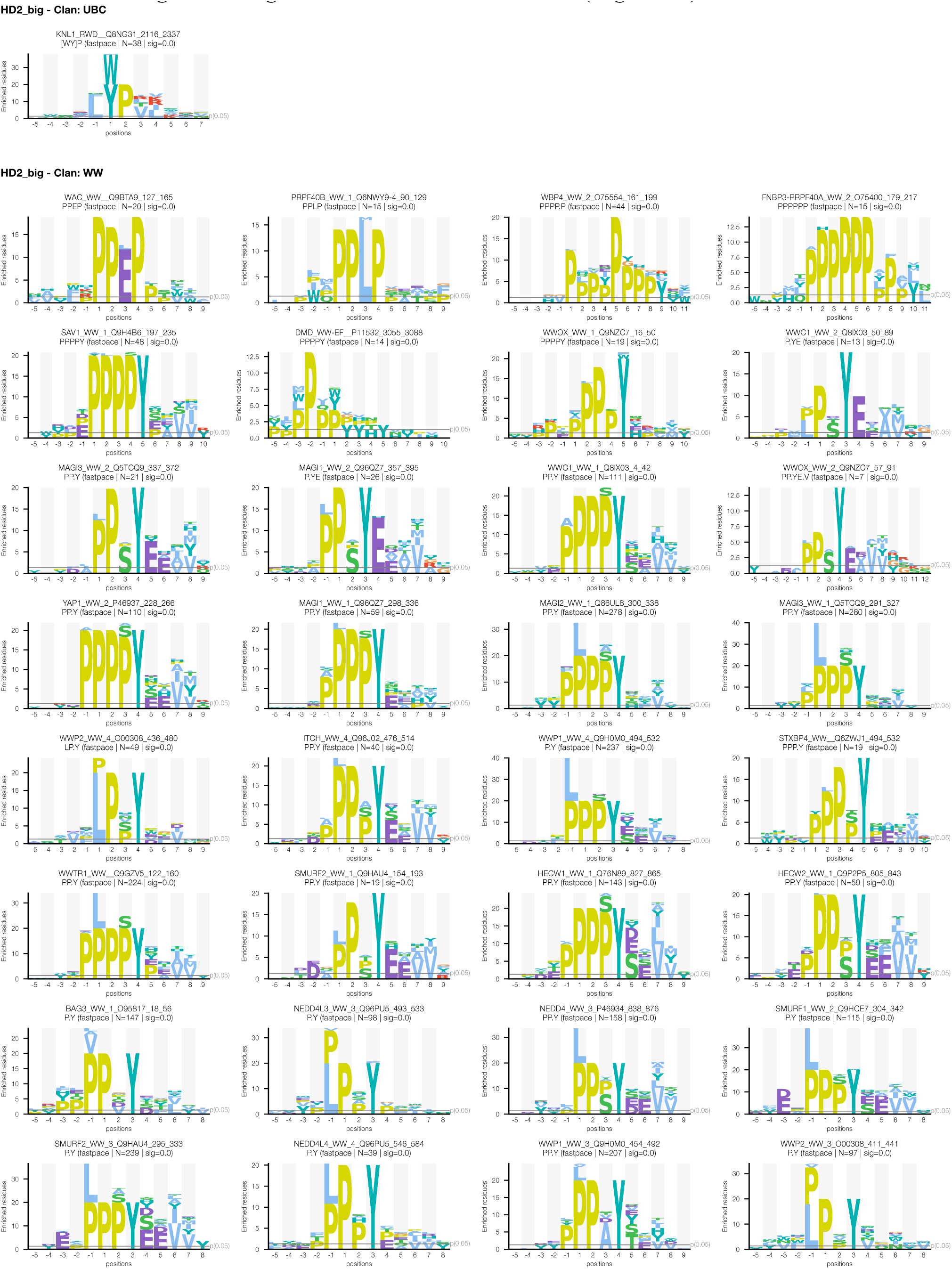
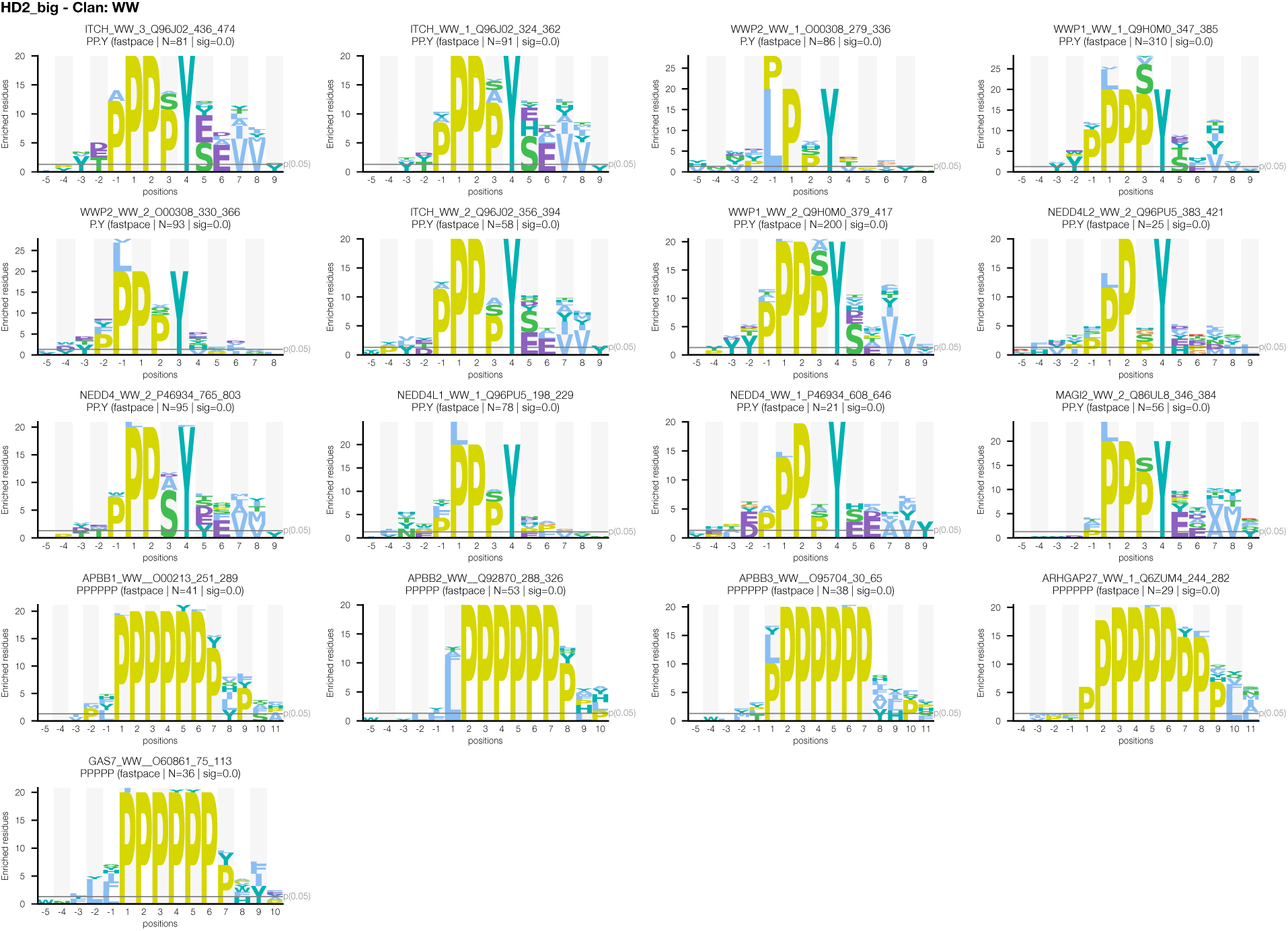
PSSMs generated based on ProP-PD data Compilation of PSSMs generated based on the ProP-PD selection results. Logo representations of the PSSMs of the motifs generated through ProP-PD selections against the HD2 library. Logos are organized following the domain clans as noted in Supplementary Table S2A. For each Logo the descriptive bait identifier is shown together with the motif’s regular expression, number of ProP-PD enriched peptides that were used for the PSSM generation (N) and the motif significance p-value (sig).

**Extended Data Figure 3.**
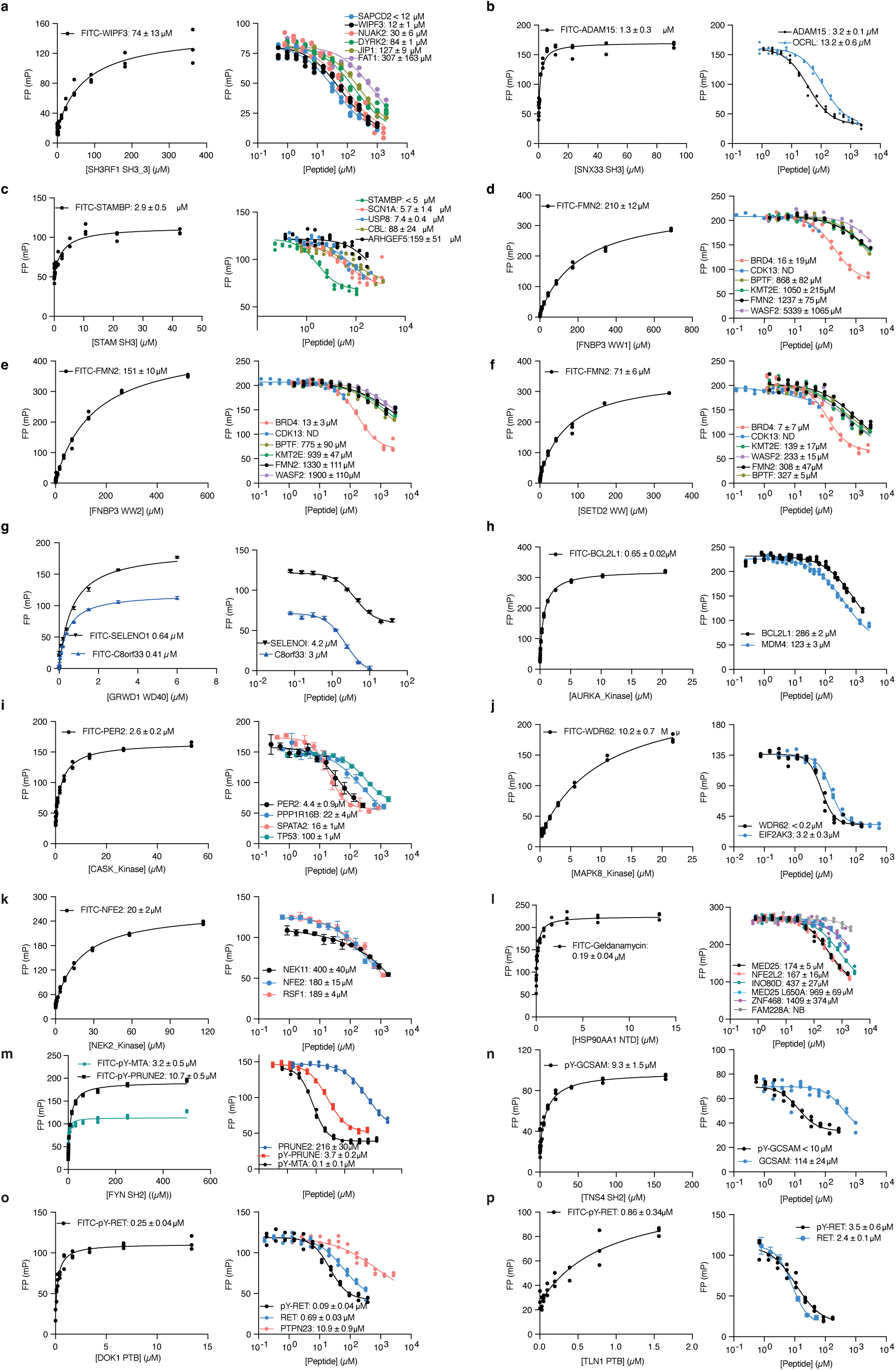
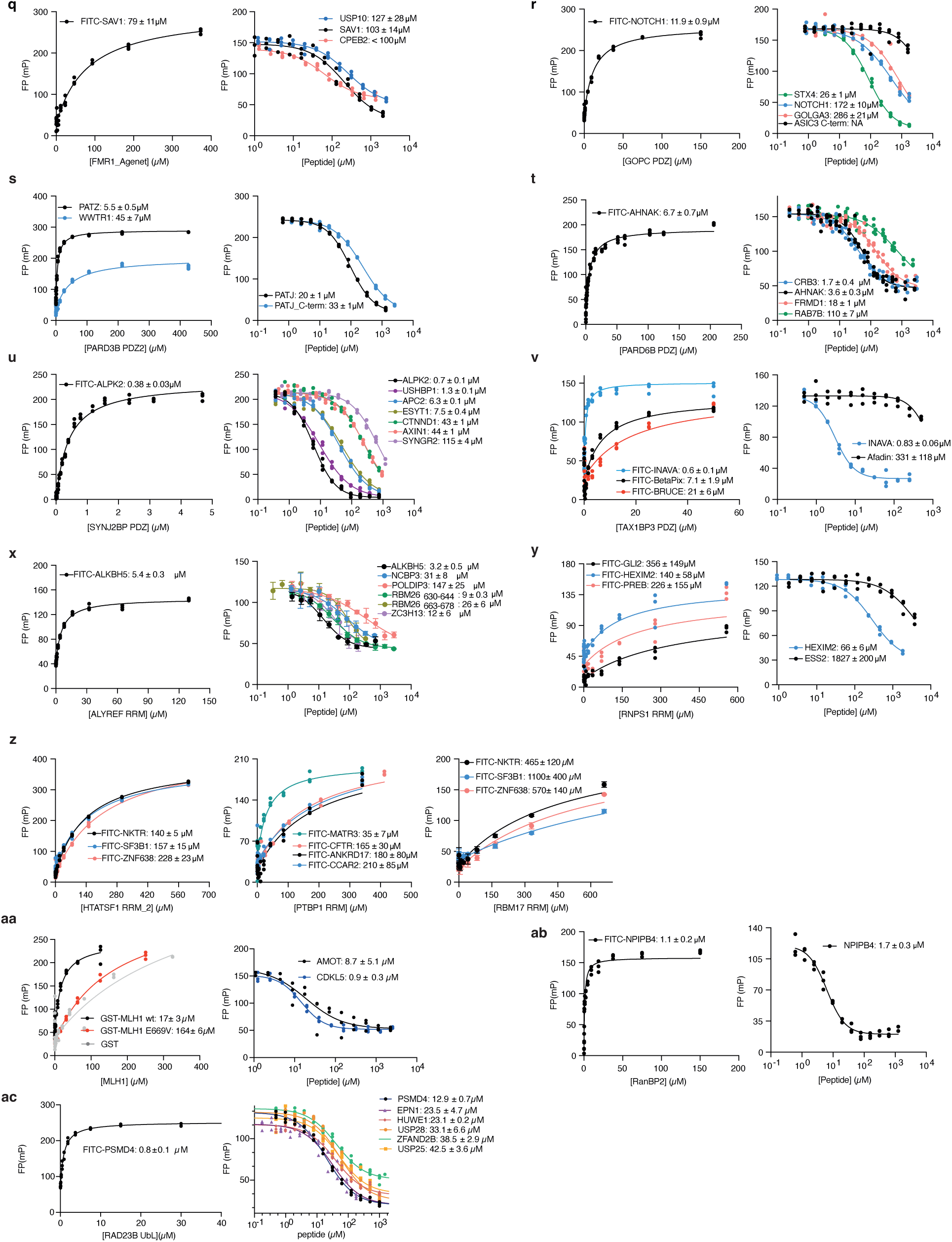
FP based affinity measurements. Shown are direct binding of the FITC-labeled probe peptides (left) and the competitive displacement of the probe by unlabeled peptides as specified in Sup. Table 3. Shown are results for **a**, SHRF1 SH3_3; **b**, SNX33 SH3; **c**, STAM SH3; **d**, FNBP3 WW1; **e**, FNBP3 WW2; **f**, HYPB (SETD2) WW; **g**, GDWR1 WD40; **h,** AURKA kinase domain; **i**, CASK kinase domain; j, MAPK8 kinase domain; **k**, NEK2 kinase domain; **l**, HSP90AA1 NTD; **m**, FYN SH2; **n**, TNS4 SH2; **o**, DOK1 PTB; **p**, TLN1 PTB; **q**, FMR1 TT; **r**, GOPC PDZ: **s**, PARD3B PDZ2; **t**, PARD6B PDZ; **u**, SYNJ2BP PDZ; **v**, TAX1BP3 PDZ; **x**, ALYREF RRM; **y**, RNPS1 RRM; **z**, HTATSF1 RRM; **aa**, PTBP1 RRM; ab, RBM17 RRM; **ac**, MLH1 NTD; **ad**, RanBP2 PH; and **ae**, RAD23B Ubl. Experiments were performed in technical repeats (n=3).

**Extended Data Figure 4.**
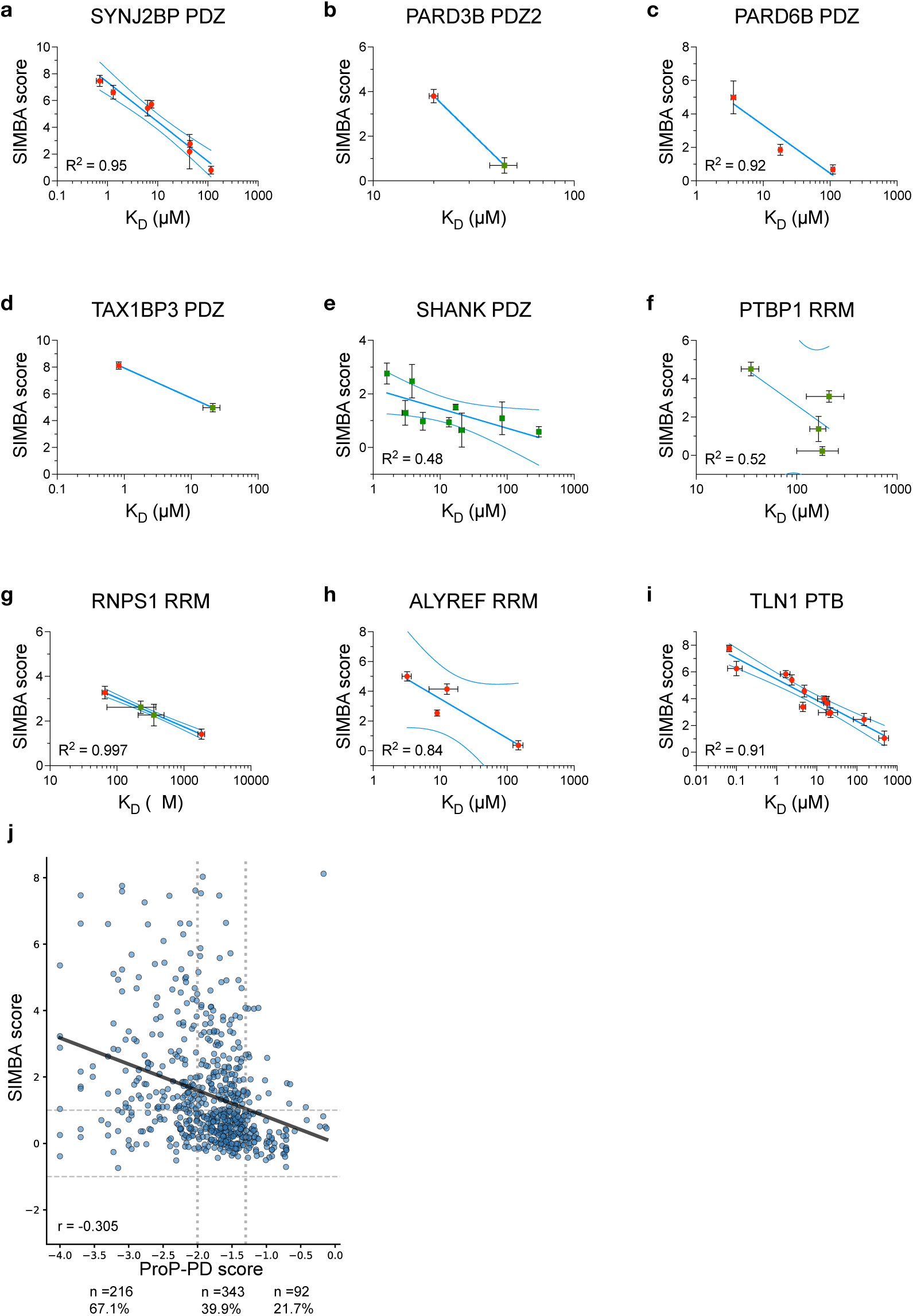
Correlation between SIMBA scores, affinities and ProP-PD scores. Plots comparing SIMBA scores with available K_D_ values for identical or overlapping peptides. Green squares denote K_D_ values obtained by direct measurement of FITC-labeled peptides; red circles denote K_D_ values obtained by competitive displacement assays; when both were available, the displacement value was used. Blue lines show the semi-log fit (solid) and 95% confidence intervals (dashed). Error bars are SD, n = 3 (K_D_) or n = 6 (SIMBA). K_D_ values are from this study or from Benz et al., 2025^40^.Shown are results for **a**, SYNJ2BP PDZ; **b**, PARD3B PDZ2; **c**, PARD6B PDZ; **d**, TAX1BP3 PDZ; **e**, SHANK PDZ; **f**, PTBP1 RRM; **g**, RNPS1 RRM; **h,** ALYREF RRM; **i**, TLN1 PTB**. j**. Correlation between SIMBA scores and ProP-PD scores.

**Extended Data Figure 5.**
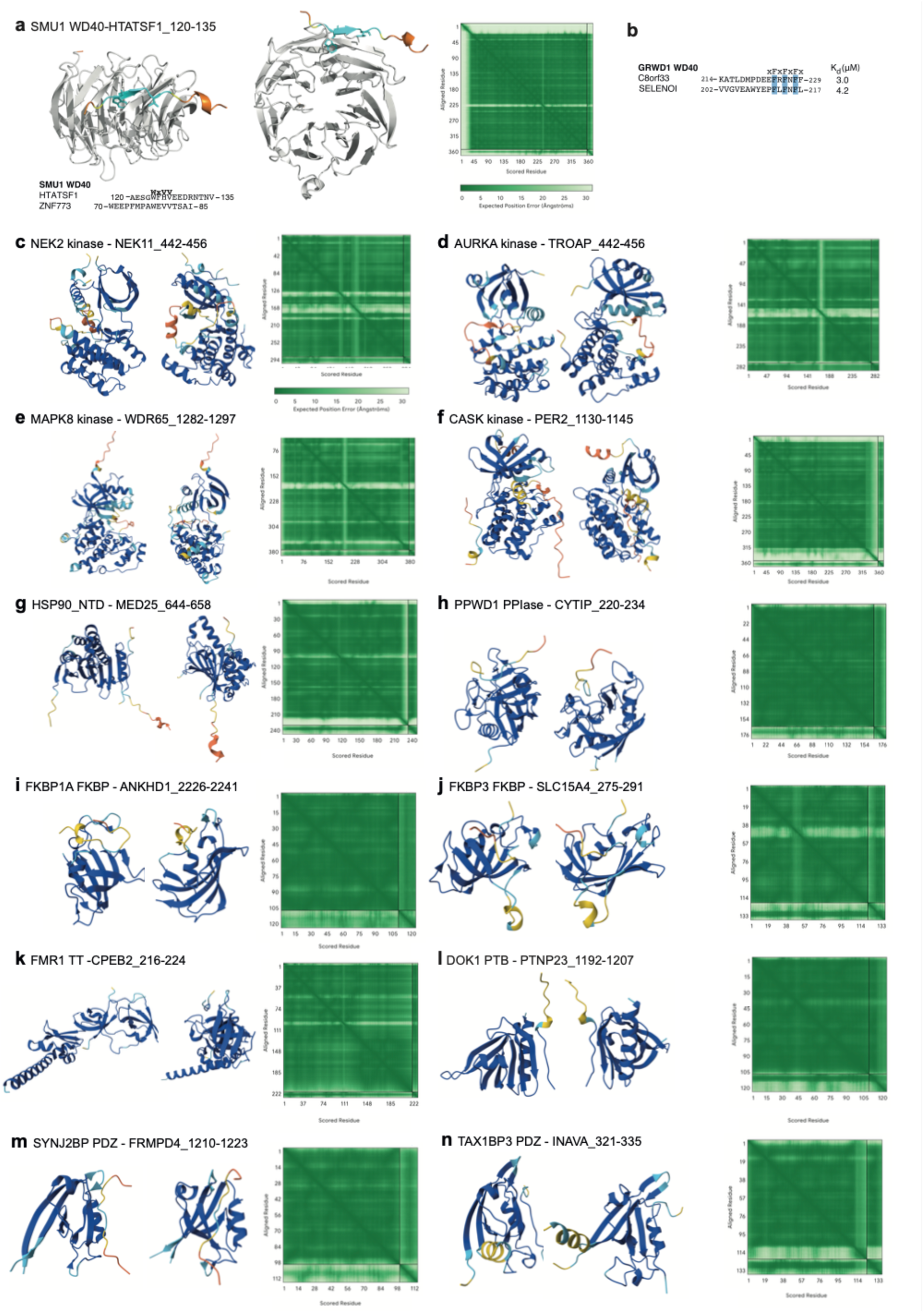
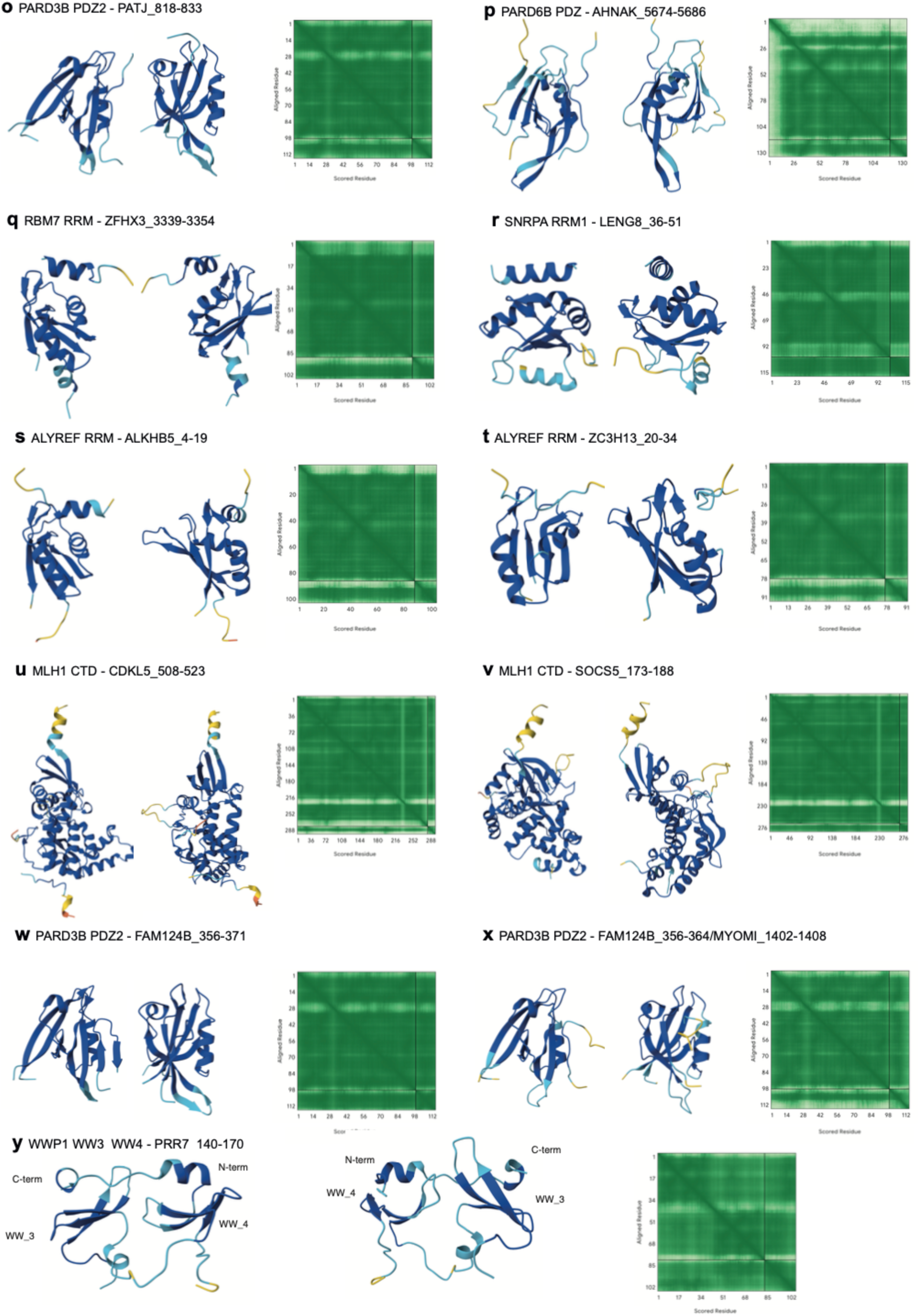
AlphaFold3 models in two representations together with the predicted aligned errors. **(a)** SMU1 WD40-HTATSF1_120-135_, (**b**) GRWD1 WD40 ligands together with determined K_d_ values, (**c**) NEK2 kinase – NEK11_442-456_, (**d**) ARUKA kinase – TROAP_442-456_, (**e**) MAPK8 kinase – WDR65_1282-1297_, (**f**) CASK kinase PER2_1140-1145,_ (g) HSP90 NTD – MED25_644-658_, (**h**) PPWD1 PPIase – CYTIP_220-234_, (**i**) FKBP1A FKBP – AHNAK1_2226-2241_, (**j**) FKBP3 FKBP – SLC15A3_275-291_, (**k**) FMR1 TT – CPEB2_216-224_, (**l**) DOK1 PTB – PTPN23_1192-1207_, (**m**) SYNJ2BP PDZ – FRMPD4_1210-1223_, (**n**) TAX1BP3 PDZ – INAVA_321-335_, (**o**) PARD3B PDZ2 – PATJ 818-833, (**p**) PARD6B PDZ – AHNAK_5674-5686_, (**q**) RBM7 RRM – ZFHX2_3339-3354_, (**r**) SNRPA RRM1 – LENG8_36-51_, (s) ALYREF RRM – ALKHB5_4-19_, (**t**) ALYREF RRM – ZC3H13_20-34_, (**u**) MLH1 CTD – CDKL5_508-523_, (**v**) MLH1 CRD SOCS5_173-188_, (**w**) PARD3B PDZ2 – FAM124B_356-371_, (**x**) PARD3B PDZ2 – FAM124B_356-364_/MYOM1_1402-1408_, (**y**) WWP1 WW3_WW4 – PRR7_140-170_. The structures are colored according the confidence of the model (blue: pIDDT >90; cyan: 90>pIDDT> 70; yellow: 70>pIDDT>50; orange < 50), with the exception for the SMU1 WD40 which is colored in grey to allow the visualization of the bound peptide ligand. The expected position error is shown to the right.

**Extended Data Figure 6.**
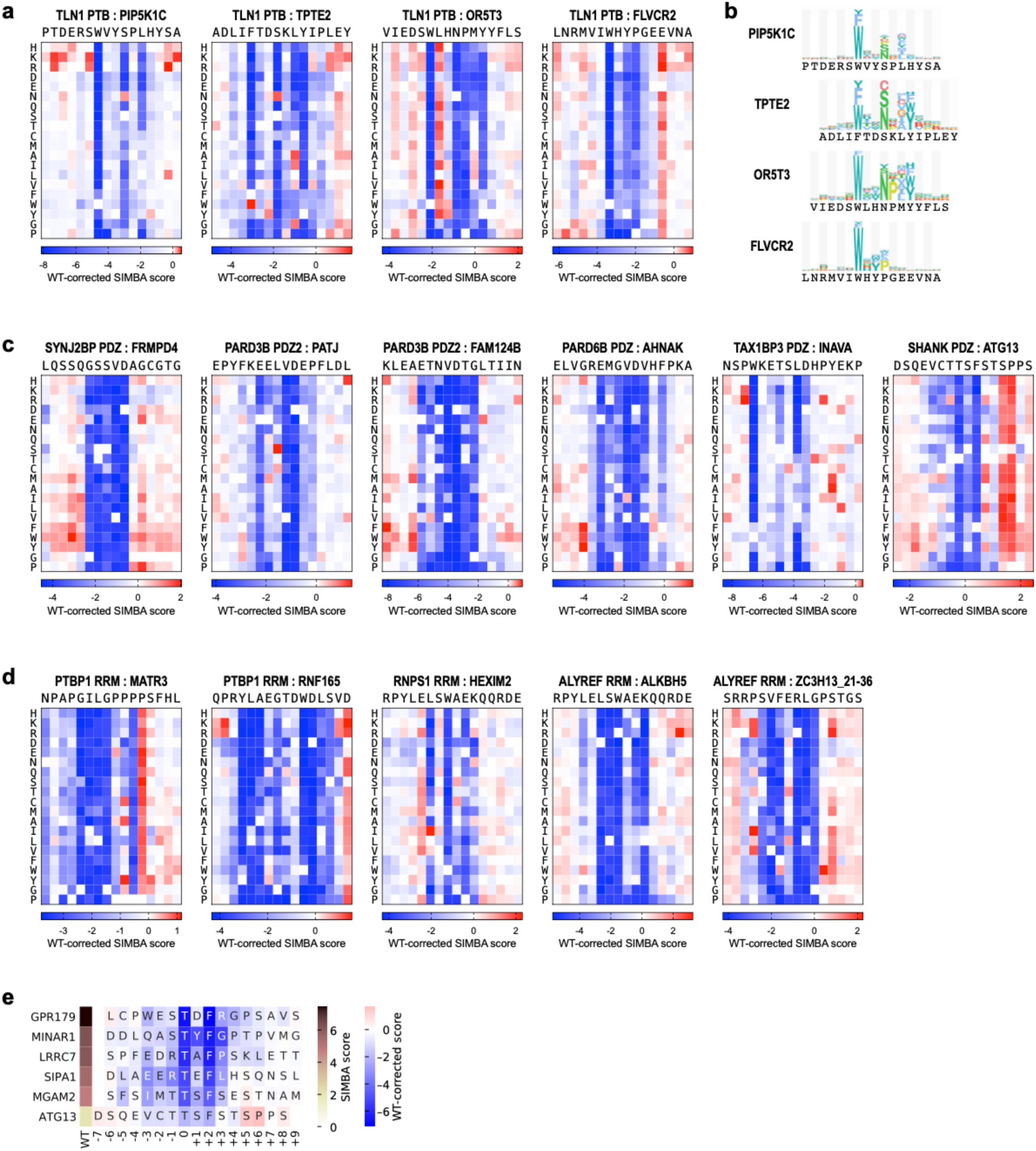
Heatmap representation of SIMBA monitored DMS analysis. **a**, **b,** DMS of four model peptides with motif variations binding to TLN1 PTB shown in heatmap representation (**a**) and as logos (**b**). **c**, DMS of peptides binding to SYNJ2BP PDZ, PARD3B PDZ2, PAR6B PDZ, TAX1BP3 PDZ and SHANK PDZ. **d**, DMS results of two peptide binding PTBP1 RRM, one peptide binding to RNPS1 RRM and two ligands of ALYREF RRM. **e**, SIMBA based alanine scanning of six SHANK1 PDZ domain binding peptides.

**Extended Data Figure 7.**
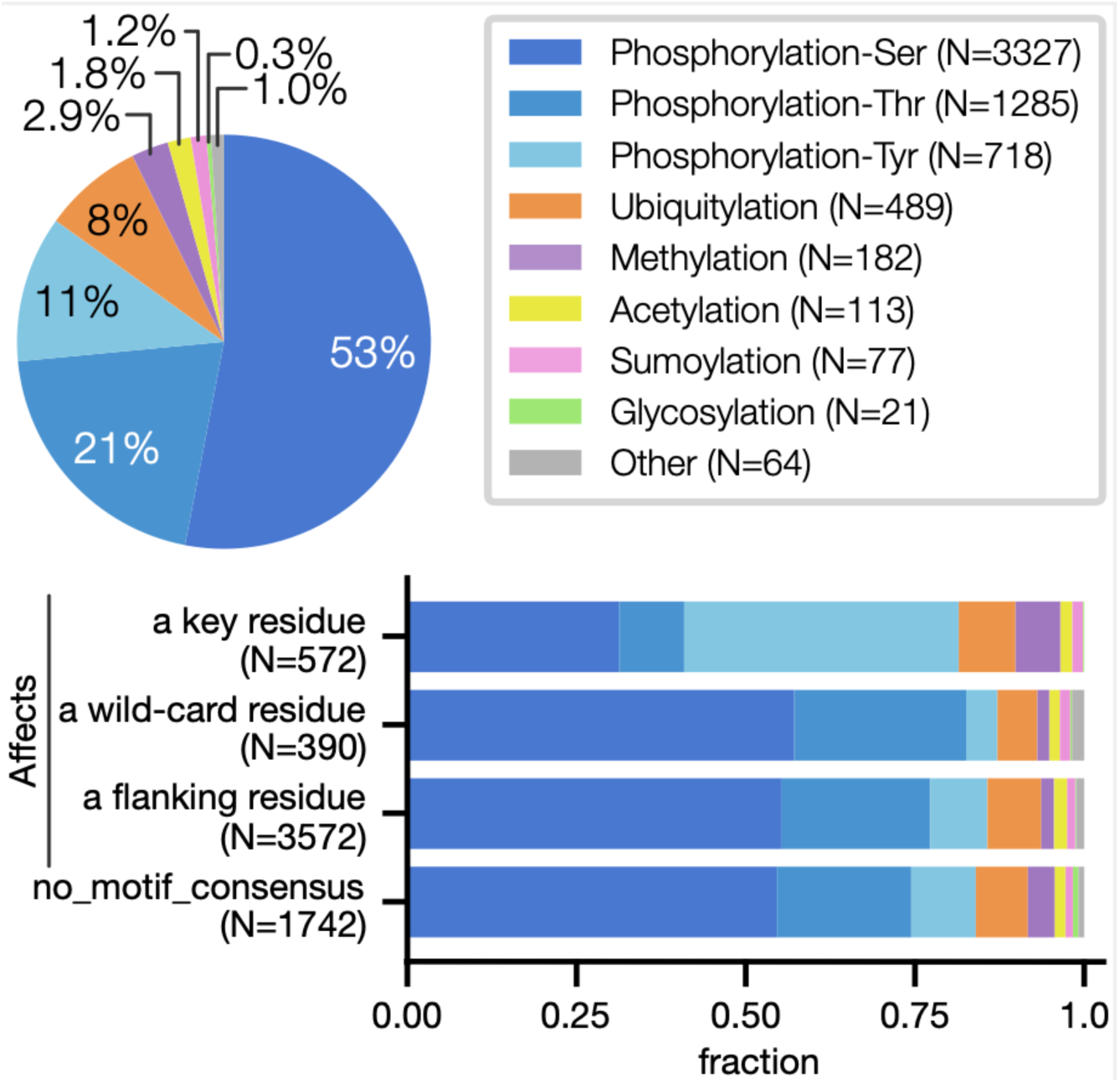
Overview of PTMs mapping to the peptides in ASHI. **Top:** Distribution of PTM types. **Bottom**: Fraction of the PTMs affecting distinct positions in the peptides in relation to the motif determinants.

## Supplementary Methods

### Protein expression and purification

Fusion proteins (**Supplemental Table 1**) were produced in *E.coli* BL21 (DE3) gold cells in 2YT (10 g yeast extract, 16 g tryptone, 5 g NaCl in 1 L) with suitable antibiotics (kanamycin 35 µg/mL, or carbencillin 100 µg/mL). Cells were grown at 37°C, 200 RPM until the OD_600_ 0.5-0.8. Protein expression was induced with 0.3 mM isopropyl β-D-1-thiogalactopyranoside (IPTG) and allowed for 20 h at 18°C. Cells were harvested and lysed in PBS (137 mM NaCl, 10 mM Na_2_HPO_4_, 1.8 mM KH_2_PO_4_ and 2.7 mM KCl), 5 mM MgCl_2_, 1 % Triton-X-100, 10 ug/mL DNase, Complete protease inhibitor cocktail (Roche) and lysozyme. Suspended cell lysate was sonicated with 70 % amplitude, 2 s pulse on and 2 s pulse off (20 s for 100 mL cultures, 5 min for 4 L cultures). Cells were incubated for 30 min at 4°C and then centrifuged for 1 h at 17,000 × g, 4 ℃. The supernatant was incubated with Glutathione Sepharose™ 4 Fast Flow (Cytiva) or Ni Sepharose excel (Sigma) for 1 h, 4°C for 1 h. The resin was pelleted by centrifugation at 500 × g and the supernatant was decanted. For GST-tagged proteins, beads were washed three times with PBS buffer and eluted with reduced glutathione in 50 mM Tris, pH 8.0. For His-tagged proteins, or His-MBP-tagged proteins, beads were washed two times (5 min each) with PBS and two times with PBS supplemented with 40 mM imidazole. Protein was eluted with 500 mM imidazole in PBS. Protein size and purity was confirmed by SDS-PAGE.

For affinity measurements, fusion proteins were captured on Ni^2+^ Sepharose excel resin. The purification tags were cleaved off using HRV-3C by on-bead cleavage. The resin with captured protein was equilibrated with buffer (50 mM Tris, 150 mM NaCl, pH 8.0, 1 mM EDTA) and incubated for overnight with HRV-3C at 4 °C. Cleaved protein was obtained by collecting the flow through. The protein was run on SDS-PAGE to confirm the cleavage. Protein was buffer exchanged with 50 mM Sodium Phosphate buffer pH 7.4 (5 mM DTT if protein has cysteines) using desalting PD-10 column. For MBP-tagged proteins, the tag was not cleaved for affinity measurements because of difficulty in cleaving the tag.

### Purification of WD40 domains

All protein constructs used for assays and structural studies were expressed in Spodoptera frugiperda (Sf9) as previously described^1^. For protein purification, cells were resuspended in the base purification buffer (20 mM Tris pH7.5, 300 mM NaCl), 2 mM TCEP, supplemented with protease inhibitor, 0.1% NP-40, and benzonase. Suspended cells were lysed by sonication and clarified by centrifugation. Supernatant was collected and incubated with cobalt resin for 1 hour, under rotation at 4 °C. Beads and supernatant were placed in an open column for affinity purification. Beads were washed with 10 to 20 column volumes (CVs) of base buffer, followed by a 5 CVs wash using base buffer supplemented with 5 mM imidazole. Protein was eluted using 5 CVs of base buffer supplemented with 250 mM imidazole. Protein was further purified by size exclusion chromatography (Superdex 200, GE Healthcare) using base buffer. In case of PPWD1, an additional ion exchange step was performed. Purification samples were analyzed by SDS-PAGE and stored at – 80°C.

### ProP-PD selections

Phage selections using the HD2 library^2^ were carried out in at least three replica per protein domains in four consecutive selection rounds. All incubation steps were conducted in the cold room (6 °C) while shaking if not specified elsewise.

#### Selection day 0

GST-tagged domains or GST control (10-15 µg) were immobilised in 96-well MaxiSorp plates (Nunc) in a total volume of 100 µL diluted in PBS and the plates incubated overnight.

#### Selection day 1

Wells were blocked with 0.5% bovine serum albumin (BSA) in PBS (200 µL/well) for 1 h. The naïve phage library (10 µL/well, 10^11^ phages) was precipitated by adding one fourth the volume of 20% PEG-8000/400 mM NaCl and incubation on ice for 15 min, followed by centrifugation. The library was resuspended in 0.5% BSA in PBS and 0.05% Tween-20 (100 µL/well). The GST coated pre-selection plates were washed four times with PBS supplemented with 0.05% Tween-20 and the naïve phage library added to each well (100 µL). Plates were incubated for 1 h. The bait coated plates were washed four times with PBS supplemented with 0.05% Tween-20 and the unbound phages transferred from the GST plates to the respective wells of the bait domain plate. After 2 h incubation, unbound phages from the bait domain plate were removed by washing five times with PBS supplemented with 0.05% Tween-20. Bound phages were eluted by adding log phase Omnimax bacteria and incubating for 30 min at 37 °C and 200 rpm (100 µL/well). Next, M13K07 helper phages (10 µL/well, 10^9^ phages) were added to each well and the plates again incubated for 45 min at 37 °C at 200 rpm. The bacteria were subsequently amplified in 2YT medium supplemented with 30 µg/mL kanamycin, 100 µg/mL carbenicillin and 0.3 mM IPTG by incubation overnight at 37 °C and 200 rpm.

#### Selection day 2-4

The steps on day 2-4 were the same as on day 1, with the exception that phages from the previous selection day were used as in-phages. For this, the phage solutions amplified overnight were cleared from bacteria by centrifugation (15 min at 1,700 xg and 4 °C). The supernatant containing the phages was pH-neutralised by the addition of 10x PBS pH 7.4 and subsequently heat-inactivated by incubation at 60 °C for 10 min. The harvested phage pools were then used as in phages for the next round of selection.

### Phage pool ELISA

Phage pool ELISA was performed to assess the enrichment of binding phages over the days of selections. For this, bait domains (7.5 µg) and GST (5 µg) were immobilised on a 384-well MaxiSorp plate (Nunc) (total of 50 µL) overnight. Plates were blocked for 1 h with 100 µL 0.5% BSA in PBS for 1 h. Phage pools from selection day 1-4 were added to both bait domain and GST wells and the plates incubated for 1.5 h. Next, plates were washed four times with PBS supplemented with 0.05% Tween-20 (100 µL/well) and bound phages were detected by the addition of anti-M13 mouse HRP-conjugated antibody (50 µL/well, 1:5000 in 0.5% BSA in PBS supplemented with 0.05% Tween-20, Sino Biological Inc). Plates were incubated for 1 h and then washed four times with PBS supplemented with 0.05% Tween-20 and once with PBS (100 µL/well). After that, 50 µL of TMB substrate (Seracare KPL) were added to each well, which was converted by the HRP if there was a bound antibody. The reaction was stopped by the addition of 0.6 M H_2_SO_4_ (50 µL/well) and the signal read-out on an iD5 (Molecular Devices) by measuring the absorption at 450 nm. Pools were considered enriched if the signal observed for the bait domain was at least two-fold higher in comparison to control wells.

### Next-generation sequencing (NGS)

The peptide coding regions of the enriched phage pool were barcoded and PCR amplified using a dual barcoding strategy^3^. For this purpose, 5 µL of each phage pool were used as template in a PCR reaction of 22 cycles using Phusion High-fidelity master mix (Thermo Fisher). PCR reactions (25 µL) were normalised by purification with 25 µL Mag-bind Total Pure NGS (Omega Bio-tek). The amplicons were pooled and further purified by gel purification in a 1% agarose gel supplemented with GelRed (1:10000) using the QIAquick Gel extraction Kit Qiagen. DNA content was quantified with Quant-iT PicoGreen dsDNA Assay Kit (Molecular probes by Life technologies) and the sample sent for NGS to the NGS-NGI SciLifeLab facility. Analysis was by Illumina MiSeq v3 run, 1 × 150 bp read setup, 20% PhiX and returned an average of 33,000 reads per set of barcodes (on average 18 million reads per NGS run). The results were analysed using in house scripts as previously described^3^. In short, NGS data was acquired in FASTQ format from where the DNA sequences were sorted based on their barcodes; sequences with an average quality of 20 were preserved and their adapters and barcodes were trimmed;

DNA sequences were translated to amino acid sequence and all sequences matching the library design were counted. The resulting data after NGS data processing consist of 16 amino acid long peptide sequences with their read counts. These tables were analysed using our previously published pipeline PepTools^2^, which maps, annotates and identifies specificity determinants (motifs) in the list of peptides. The annotated peptides were then categorized as described before^2^ into “confidence levels” based on four confidence metrics: NGS counts, presence of the same peptide in replicate selections, presence of overlapping peptides and matching of the specificity determinants identified for the bait (if any); depending on how many of the metrics were met, each peptide was given a score between 0 and 4, with a confidence level of 4 being considered “high” and a confidence level between 2 and 3 considered as “medium”. The specificity of each peptide was also calculated as the proportion of the normalized counts for that peptide in the set of selections with each specific bait domain compared to all other domains where the peptide was enriched. This value allows us to identify promiscuous peptides. The scoring according to the established protocol is provided for legacy reasons^2^, and used together with an optimized scoring protocol.

### Optimizing the ProP-PD peptide data integration and scoring

A composite ProP-PD score integrating multiple peptide features was developed to identify high-confidence bait-interacting peptides. For each peptide returned from each bait screen, six features were extracted: (i) replicate count, the number of independent biological replicates in which the peptide was enriched; (ii) overlap count, the number of overlapping peptides within the phage display library that were co-enriched; (iii) mean proportion of NGS counts, the mean fraction of next-generation sequencing reads attributed to the peptide relative to all peptides for the bait; (iv) motif consensus enrichment p-value, a statistical measure of enrichment of the peptide sequence by a position-specific scoring matrix derived from a FaSTPACE analysis^4^ of the replicated and overlapping peptides; (v) peptide-bait specificity, the proportion of NGS counts for the peptide that are observed for the given bait; and (vi) peptide-bait promiscuity, the number of baits that returned the peptide. To account for bait-specific variation in screening depth and coverage, replicate and overlap counts were independently normalised within each bait protein by log_10_-transformation relative to the maximum observed value for that bait. All continuous features were z-score standardised (mean-centred and scaled to unit variance) across the complete dataset prior to model training.

Peptide candidates were labelled as one of two classes: curated motifs, comprising peptides with validated experimental or structural evidence of interaction between the bait protein and the peptide-containing region of the prey protein as annotated in MoMaP (https://slim-tools.org/momap/), and other peptides, comprising all remaining candidates lacking sufficient supporting evidence of bait interaction. Model training was restricted to bait proteins containing at least one curated motif, ensuring that only high-quality, experimentally grounded ground truth labels informed the classifier.

A supervised partial least squares (PLS) regression model was trained to integrate the six peptide features into a unified continuous interaction score. A single-component PLS model (n_components = 1) was fitted to binary target labels (curated motif = 1, other = 0), with the first latent component used directly as a continuous interaction score. Model performance was assessed by stratified k-fold cross-validation (k = 5). Stratification ensured that the proportion of curated motifs relative to other peptides was preserved within each fold, preventing class imbalance. The cross-validation procedure was as follows: for each fold, all features in the training partition were independently standardised using z-score normalisation. A single-component PLS model was then fitted to the standardised training features and binary target labels. Continuous scores generated on the held-out test partition were interpreted as probabilistic predictions of bait interaction propensity, with higher scores indicating greater likelihood of a curated motif classification. ROC and precision-recall (PR) curves were computed for each fold, and area under the ROC curve (ROC-AUC) and area under the PR curve (PR-AUC) were calculated per fold, yielding five independent performance estimates that were subsequently averaged to produce overall model performance metrics. To assess the contribution of individual features, per-feature ROC-AUC values were computed independently across all folds by scoring each feature in isolation, enabling ranking of feature importance within the multivariate framework.

Following cross-validation, a final PLS model was trained on the complete curated training dataset, comprising all bait proteins with at least one curated motif, using the same standardisation and fitting procedure described above. The first latent component of this final model was applied to the entire peptide dataset, including peptides from baits not represented in the training set, to generate a continuous supervised PLS score for every peptide. Higher ProP-PD supervised PLS scores indicate greater predicted likelihood of peptide-bait interaction, enabling quantitative ranking of all candidate peptides for downstream experimental validation.

Logos were built for baits where FaSTPACE or SLiMFinder identified a motif de novo. For this all peptides annotated as motif containing were aligned centered on the motif and the binomial log probability was calculated for each amino acid at each position of the aligned motifs. The expected frequency for each amino acid was calculated from the human disorderome (defined as the regions with a IUPred disorder cut-off of 0.5). The plotting of the logos was performed using Python’s library Logomaker^5^.

To establish an operational classification threshold, ROC analysis was performed by comparing supervised PLS scores against the curated motif dataset. The optimal threshold was identified as the score value that maximised Youden’s J statistic (J = sensitivity + specificity − 1), which balances sensitivity and specificity simultaneously. To support reporting at multiple stringency levels, false positive rates (FPR) and true positive rates (TPR) were additionally computed at three predetermined FPR thresholds (FPR ≤ 0.05, 0.01, and 0.001), providing a standardised framework for selecting peptide sets of varying confidence for experimental follow-up. ProP-PD scores were defined as log10 of the FPR. ProP-PD scores were benchmarked against ProP-PD confidence scores from the original ProP-PD bioinformatic prediction pipeline by computing ROC-AUC and PR-AUC for both scoring methods against the curated motif dataset, enabling direct quantitative comparison of discriminative performance (**Supplementary Figure 9**). The log10 of the FPR provides a continuous scoring, we provide confidence bins of medium (−1.3), high (−2), and excellent (−3)- The results of the currently and previously generated data was processed following the optimized protocol (**Supplemental Table 2**).

### Co-occurrence enrichment analysis

The enrichment of bait-bait domain pairs from different domain clans binding to same sets of motif-containing proteins was performed as described before^6^. In short: For each bait-bait pair of different domain clans that showed binding to co-occurring motif containing proteins, an enrichment p-value was calculated by comparing the observed shared prey proteins against randomized samples of peptides matching the enrichment results for each bait. The randomized samples were generated from the set of enriched peptides for all baits. 1,000,000 randomized sets were generated and used to calculate the probability of the observed overlap. Only peptides with a PLS score of high or excellent were considered for the analysis.

### Fluorescence polarization experiments

For FP experiments, FITC-labelled peptides were dissolved in DMSO and unlabeled peptides in 50 mM sodium phosphate pH 7.4. Black non-binding 96-well plates (Corning) were used for the measurements, and the read-out was on an iD5 plate reader (Molecular Systems) by exciting the FITC-label at 485 nm and measuring the emitted light at 535 nm in two different angles. From this, the FP signal is calculated in mP. For saturation experiments, a 1:1 dilution series of protein is generated (25 µL), to which 25 µL of 10 nM FITC-labelled peptide (diluted in 50 mM sodium phosphate pH 7.4) are added to an end volume of 50 µL in each well. In case of the displacement experiments, a pre-complex of protein (concentration = 1-2 fold K_d_) and FITC-labelled peptide (10 nM) is prepared in 50 mM sodium phosphate pH 7.4. The unlabeled peptide (25 µL) is then titrated in a 1:1 dilution series against the complex (25 µL), so that the end volume is again 50 µL in each well. The read-out on the plate reader is immediate after pipetting the plates. For proteins or peptides containing cysteine residues, 5 mM DTT was added in the buffer.

FP binding assays for the WD40 domains were performed following a slightly modified protocol, using 20 mM Tris pH 7.5, 150 mM NaCl, 5% glycerol, 0.5 mM TCEP, 0.01% Triton buffer. All tested proteins were titrated from an initial 4 -40 µM highest concentration, followed by a factor 2 serial dilution. FITC peptides binding to proteins were tested at 20 nM final concentration. The assay was incubated for 1 hour at room temperature and plates were read using BioTek Synergy 4 (BioTek) plate reader. Prior to displacement assay, proteins at concentration corresponding to 80% maximum FP signal were mixed with FITC peptides at 20 nM final concentration. Displacement was evaluated by using a titration curve of unlabeled peptide, with 40 µM as highest concentration. Buffer, incubation time and reading conditions followed FP binding protocol. All FP analyses were performed in triplicates and, at least, two independent experiments. All K_d_ and K_d_^DISP^ values were calculated using GraphPad Prism version 10.0.

### SPR

MLH1 constructs including wt, E669V MLH1 and L607R MLH1 were cloned in pETM11 vector using NcoI and EcoRI restriction sites. Proteins were expressed and purified as described in section Protein Expression and Purification. Proteins and peptides were suspended in 50 mM Sodium Phosphate pH 7.4, 5 mM DTT. Surface Plasmon Resonance (SPR) experiment was performed on Biacore 3000 (Uppsala University) instrument using NTA sensor chip (BR100407, Cytiva). The sensor surface was activated with 1:1 mixture of NHS/EDC (Cytiva) at a flow rate of 10 μL/min for 7 min. This was followed by injection of 500 mM Nickel chloride at a flow rate of 10 μL/min for 1 min, respectively. His-tagged proteins at concentrations 0.1 mg/mL was injected in 50 mM Sodium Phosphate pH 7.4, 5 mM DTT buffer at a flow rate of 2 μL/min for 10 min. Then surface was deactivated using ethanolamine at a flow rate of 10 μL/min for 7 min. Proteins were immobilized at 2500 RU. Peptides, CDKL5 and SOCS5 were prepared as a series of five two-fold serial dilutions in the same running buffer, 50 mM Sodium Phosphate pH 7.4, 5 mM DTT. Stock solutions of CDKL5 and SOCS5 were 200 μM and 147 μM, respectively. Peptides were injected over the immobilized protein at a flow rate of 30 µL/min with a contact time of 60 s, followed by dissociation time of 120 s. Sensograms were collected and analyzed using BIAevaluation software. Analyzed data was exported and plotted using Graphpad prism 10.

### Peptide SPOT-array analysis

Peptides SPOT-arrays were purchased from JPT (PepSpots). The peptide sequences designed were flanked with one glycine residue both at N-terminal and C-terminal. Purchased membranes were activated with methanol by incubating for 5 min and then washed three times (3 min each) with TBST (20 mM Tris, 150 mM NaCl pH 8.0, 0.05 % Tween-20). Membranes were blocked in TBST with 3 % BSA for 1 h at 4 ℃. The purified proteins were buffer exchanged with a blocking buffer having 1 mM

DTT. Membranes were incubated with GST-tagged or His-tagged protein overnight at 4 ℃. Washed the membranes three times (1 min each) with TBST buffer. Membranes were incubated with HRP-conjugated antibody (anti-GST: GERPN1236 Sigma or anti-His: MA1-21315-HRP Thermo fisher) for 1 h at 4 ℃. Membranes were washed again three times (1 min each) with TBST buffer followed by a wash with TBS buffer. The arrays were imaged on ChemiDoc Imaging system (Bio-Rad) using ECL reagent (Clarity Max Western ECL substrate, 1705062, Bio-Rad).

### Co-immunoprecipitation (Co-IP)

For Co-IP experiments, HEK293T cells were cultured in Dulbecco’s Modified Eagle Medium (DMEM) (Thermofisher) supplemented with 10 % (v/v) fetal bovine serum (FBS) (fisher scientific) and 1 % penicillin streptomycin (Fisher scientific) at 37 °C, 5 % CO_2_ in humidified environment. Cells were seeded with 30 % confluency in 175 cm^2^ flasks. After 24 h, transfection mixture was prepared by adding either GFP control plasmid or GFP-tagged plasmids and FLAG-tagged plasmids in JetOptimus buffer, (100 µL/µg plasmid) and the JetOptimus reagent was added to the mixture (1 µL/µg plasmid). Transfection mixture was incubated for 15 min at room temperature. The mixture was added to the cells and cells were incubated at 37 °C, 5 % CO_2_. Media for cells was changed after 18 h. After 48 h from the transfection, cells were harvested. Cells were lysed with 600 µL of lysis buffer having 50 mM NaCl, 50 mM Tris, 1 mM EDTA pH 7.4, 0.1 % NP-40 (Igepal), 1 mM DTT, complete EDTA free protease inhibitors, phos-stop phosphatase inhibitors. Cells were incubated with lysis buffer for 1 h at 4 °C, followed by centrifuging at 13,600 RPM, 45 min. Supernatant was collected in fresh tubes and the total protein concentration was calculated using Bradford (BSA as standard). Protein concentrations were normalized. 100 μg of the sample was aliquoted and stored at −20 °C as input samples. Each sample was incubated with 20 μL of GFP beads for 1 h at 4 °C. Beads were washed three times with 1 mL of wash buffer having150 mM NaCl, 50 mM Tris pH 7.4, 0.05 % NP-40 (Igepal), 5 % glycerol, 1 mM DTT. Beads were suspended with 20 μL of twofold NuPAGE LDS Sample buffer (Novex). Samples were run on 4-12 % BIS-Tris NuPAGE (Invitrogen) using MOPS (Invitrogen) as running buffer. Samples were transferred on Trans-Blot Turbo Mini 0.2 µm Nitrocellulose membrane (Bio-Rad) using Trans-Blot Turbo Transfer System (Bio-Rad). Membrane was blocked in PBST 5 % skimmed milk for 1 h at room temperature. Primary antibodies (anti-FLAG M2 in mouse from Merck and anti-GFP in rabbit from Proteintech) were diluted 1:5000 in PBST 2.5 % skimmed milk. Membrane was incubated with primary antibody at 4 °C for overnight. Membrane was washed three time for 5 minutes each with PBST. Secondary antibodies ( IRDye® 800CW Goat anti-Rabbit IgG Secondary Antibody and IRDye® 680RD Goat anti-Mouse IgG Secondary Antibody from LICORbio) were diluted 1:5000 in PBST 2.5 % skimmed milk. Incubated the membrane with secondary antibody for 1 h at room temperature. Washed the membrane again three time for 5 minutes with PBST. Blots were imaged on Odyssey CLx (LI-COR).

### Systematic Intracellular Motif Binding Analysis (SIMBA)

#### Library design

Libraries for SIMBA screens encoded 16-mer peptides, flanked by flexible linkers (GSGG), inserted into a plasmid encoding a yeast signaling protein chimera Ste20^Ste5PM^ ^7,8^. Two distinct libraries were tested independently. Library 1 encoded peptides designed to test binding to nine domains (5 PDZ, 3 RRM, 1 PTB) and included a mixture of ProP-PD hits, candidate binders identified via SLiMSearch, plus mutational scanning of parent peptides. Library 2 encoded additional peptides designed for four of the original nine domains (2 PDZ, 1 RRM, 1 PTB), including mutational scanning plus chimeric peptides that swap core and flanking sequences from PARD3B^PDZ^ binders. Mutational scanning involved both saturation mutagenesis and alanine-scanning. Saturation mutagenesis tested variants with all 20 amino acids at each position in the 16-mer. Alanine-scanning replaced each position in the 16-mer individually with alanine or with glycine when the parental residue was alanine. In total, the two libraries incorporated saturation mutagenesis of 15 peptides and alanine-scanning of 15 peptides. In addition to the experimental peptides, both libraries shared a set of 55 control peptides: 25 LP motifs (that bind yeast Cln2), 7 sequences containing STOP codons, 13 NLS peptides, and 10 mutationally-inactivated NLS peptides. Library 2 included an additional set of 50 negative control peptides representing 5 inactivating point mutations in each of 10 peptides screened in library 1. All peptides in each library were represented by 3 synonymous nucleotide sequences, and each library was screened in 2 independent biological replicate experiments, providing 6 independent measurements for each peptide (2 replicates each of 3 synonymous nucleotide sequences).

#### Construction of plasmids harboring bait domains and peptides for SIMBA

The reading frames of peptide-binding “bait” domains were cloned between GST and yeast *CLN2* in a *P_GAL1_-GST-CLN2* expression cassette, pPP3572^9^, harbored in a low copy number (CEN/ARS) plasmid with a *HIS3* selectable marker. Peptide-encoding sequences were inserted into a site in the N-terminal region of a Ste20^Ste5PM^ chimera harbored in a low copy number (CEN/ARS) plasmid with a *URA3* selectable marker. The 16-mer peptide sequences flanked by flexible linker sequences (Gly Gly Ser Gly) were encoded by oligonucleotide pools (GenScript), which were amplified by PCR (12 cycles) using primers that anneal to the common flanking linkers and include AatII and NheI restriction sites. The PCR products were digested with AatII and NheI, treated with calf intestinal phosphatase, and then ligated into the AatII/NheI-digested Ste20^Ste5PM^ recipient plasmid, pPP4375^7^. The ligation products were transformed into *E. coli* (XL-10 Gold Ultracompetent Cells; Agilent/Agilent Technologies) and plated on LB+Amp plates. To obtain a good representation of the designed library, at least 30-fold greater number of bacterial transformants compared to the number of unique nucleotide sequences in the library were harvested and used for plasmid preparation from the pool of colonies. The plasmids used in this study are listed in Supplementary Table 9.

#### Yeast competitive growth experiments

All yeast experiments used strain PPY2617^7^, in the common laboratory strain background W303. Standard methods were used for growth and genetic engineering of yeast. Yeast cultures were grown at 30°C in yeast extract/peptone medium with 2% glucose (YPD) or in synthetic complete medium (SC) lacking uracil and histidine supplemented with 2% glucose or raffinose. To conduct competitive growth assays, strain PPY2617 was co-transformed with the *URA3*-marked Ste20^Ste5PM^-peptide plasmid library and one of the *HIS3*-marked bait constructs. The transformation mixture was plated on SC medium plates lacking uracil and histidine, and then incubated at 30°C for three days. Diluted aliquots from the transformation were used for colony counting to ensure that the number of transformants exceeds the number of nucleotide sequences in the library by at least 10-fold. The transformed cells were collected from the plates, diluted to 50 ml SC-URA containing raffinose and grown at 30°C for 4-8 hours. Then, the 50 ml cultures were diluted to OD_660_ 0.05 and grown overnight. At OD 0.8-1, expression of the bait domain-Cln2 fusion was induced by adding 2% galactose. After 75 minutes, an aliquot of each culture (40 mL; ∼ 3 × 10^8^ cells) was collected by centrifugation and flash frozen in liquid nitrogen to obtain the t0 sample. At the same time, yeast mating pheromone (α-factor) at 500 nM final concentration was added to the culture to begin the competitive growth. The cultures were grown at 30°C and diluted every 12 hours to keep the OD_660_ below 1. Aliquots (20 mL; ∼ 3 × 10^8^ cells) were harvested at 20 and 32 h after α-factor addition. Harvested cells were, washed with water, transferred to 1.5 ml tubes, and centrifuged again. The cell pellets were frozen in liquid nitrogen and stored at −80°C.

#### Sample preparation for deep sequencing

DNA was purified from yeast cells using the Zymo Research ZR Plasmid Miniprep Kit (#D4015). Frozen cell pellets were thawed and suspended in 200 μL of solution P1, and then were lysed using Zymolyase (0.2 units/μL; Zymo Research #E1005) for 1.5 h at 37°C, before proceeding with the remaining purification steps. Samples of plasmid DNA (4 μL) were subjected to PCR (17 cycles, 50 μL total volume) with primers that include standard P5 and P7 sequences for binding to Illumina flow cells during NGS (Primers in Supplementary Table 9). The forward primer included a P5 sequence followed by an Illumina sequencing primer binding site, a 6-nucleotide bar code, and a plasmid-annealing sequence upstream of the peptide-encoding insert; the reverse primer included a P7 sequence followed by a 6-nucleotide i7-index sequence, an i7 sequencing primer binding site, and a plasmid-annealing sequence downstream of the peptide-encoding insert. Aliquots (5 μL) of the PCR products were run in 1.2% agarose gels to confirm the presence of the desired product, and the remainders were purified using Zymo Spin I columns (Zymo Research #C1003-250) and eluted in 10 mM Tris-HCl, pH 8. The concentration of the eluted products was measured using nanodrop, and a mixture was prepared containing equal amounts of each DNA product. The PCR amplicon libraries were sequenced at Novogene.

#### SIMBA data processing

2FAST2Q^10^ was used to obtain the counts of different peptide-encoding oligonucleotide sequences from the deep sequencing data. To calculate enrichment scores, the following data processing steps were performed. (1) Oligonucleotides with <20 reads at t0 were dropped from further analysis. (2) The frequency of each oligonucleotide within a sample at t0, t20 and t32 was calculated. (3) The log2 fold change of t20 and t32 frequencies compared to the t0 frequency was calculated, and then divided by the timepoint in hours, resulting in a log2 slope value. (4) The log2 slope values were averaged for t20 and t32. (5) For each bait experiment, the average log2 slope values were z-normalized using the mean and standard deviation of a set of negative controls; for library 1, these were the peptides designed for unrelated domains (e.g., for PDZ baits, the peptides designed for RRM and PTB domains were used as negative controls), whereas for library 2 these were the set of 50 peptides included as negative controls. (6) For each peptide, the median z-score in the unfused Cln2 strain was subtracted from the median z-score in each bait domain Cln2 fusion strain to obtain the final SIMBA scores. These median values were derived from 6 measurements (3 synonymous oligos in 2 replicate experiments).

Statistical tests were performed only for SIMBA scores from designed, native peptides (i.e., excluding mutant variants). P-values were calculated using Student’s two-tailed t-test, comparing the set of scores for each peptide in the bait strain vs. the unfused Cln2 strain. Then, adjusted p-values were calculated using Benjamini-Hochberg correction via the Python package statsmodels.stats.multitest (fdr_bh), and was performed independently for each bait strain.

To create logos of sequence preferences, SIMBA scores were transformed into a preference PSSM as described previously^7^. First, the score for each amino acid variant was normalized to the lowest and highest scores in a given motif array. Then, these normalized scores were converted to a frequency metric by dividing each by the sum of all scores at the same position. Finally, the frequency metric was converted to a preference metric by subtracting 0.05, so that a neutral preference is represented by zero, favored residues are positive, and disfavored residues are negative. These preference scores were used to generate sequence logos, compare predicted versus observed binding strength, SIMBA scores were normalized as described above and then transformed into a difference PSSM^7^ by subtracting the average normalized score of all residues at a given position from the value of each residue at that position: (difference score) = (residue score) – (position average). Then, to obtain a prediction for a given peptide, the corresponding PSSM values for each residue at each position was summed across all motif positions to calculate the predicted score (PSSM sum).

### NMR

#### Peptide synthesis

The *LLN-containing peptide* (PQLRSLLLNPPPPQTY) was synthesized on a PeptiPilot (PeptiSystems®) automated peptide synthesizer, employing flow-through solid-phase peptide synthesis (SPPS) technology. The synthesis was performed at a 0.616 mmol scale using Fmoc-Rink Amide AM resin (loading: 0.410 mmol/g). In each coupling cycle, a pre-activated mixture of the incoming Fmoc-amino acid (2.5 equiv), Oxyma Pure (2.5 equiv), and DIC (5.0 equiv) in DMF was prepared and allowed to pre-activate for 5 min. The solution was then pumped through the column to remove the DMF in the column and recirculated at 50 °C for 60 min to effect coupling. No intermediate wash was performed after the coupling step. Following coupling, a capping solution (0.3 M acetic anhydride and 0.3 M pyridine in DMF) was passed through the column (1 column volume) for 5 min. To ensure complete delivery of the capping reagent, 1.3 column volume of DMF was used as a push volume, followed by a 2 column volumes DMF wash. For Fmoc deprotection, a solution of 20% (v/v) piperidine in DMF was pumped through the column. The progress of deprotection was monitored in real-time by UV absorbance at 294 nm, which detects the dibenzofulvene-piperidine adduct (DBF). The deprotection flow was halted once the UV signal dropped below 40 mAU, indicating completion. The resin was then washed with 4 column volume of DMF. Upon completion of these steps—coupling, capping, deprotection, and washing—one synthesis cycle was finished, and the instrument proceeded to the next cycle. Subsequent to the synthesis, 120 mg of the resin-bound *LLN-containing peptide* was cleaved using a 95:2.5:2.5 mixture of TFA:TIPS:H_2_O (2 mL) for 150 min and evacuated into ice-cold ether (30 mL). A further 1 mL of the cleavage cocktail was added to the resin for a further 30 min and subsequently evacuated into the ether. The suspension was centrifuged at 4000 rpm for 5 min. The supernatant was decanted, and the pellet resuspended in ice-cold ether (30 mL). This process was repeated for a total of three times, after which the peptide was dissolved in a minimal amount of water and lyophilised to give the crude peptide. The peptide was purified by preparative HPLC, using a reverse phase Kinetix C8 column (5 μm, 250 x 21.2 mm) on a VWR LaPrep HPLC system. The peptide was eluted over a 5-35% gradient (acetonitrile in H_2_O; 0.1% formic acid, detecting at 214 nm) over 26 min at a flow rate of 10 mL min-1. Following lyophilisation, the peptide was acquired as a white powder. Purity was determined by analytical HPLC, using a Kinetix C8 column (5 μm, 150 x 3.0 mm) on a VWR LaChrom Elite HPLC system. The peptide was eluted over 40 min (acetonitrile in H_2_O; 0.1% formic acid, detecting at 214 nm) at a flow rate of 0.5 mL min-1, and determined to have >95% optical purity. Expected [M+2H]^2+^: 917.0; observed 917.3; Expected [M+3H]^3+^: 611.7; observed 611.9.

#### Protein expression and purification

HSP90 (P07900) N-terminal domain (aa 9-236)^11^ encoded in a pGEX-4T3 vector (Addgene plasmid # 22481) and transformed in E.coli BL21(DE3) gold bacteria (Agilent Technology). The bacteria were grown 8 x 750mL 2YT (10 g yeast extract, 16 g tryptone, 5 g NaCl per L). When reaching an OD600 of 0.6, protein expression was induced with 1 mM isopropyl β-D-1-thiogalactopyranoside and was allowed to proceed for 20 h at 18°C. Bacteria was harvested for 10 min at 4,500 xg. The bacterial pellet was resuspended in lysis buffer (75 mL PBS (137 mM NaCl, 2.7 mM KCl, 95 mM Na_2_HPO_4_, 15 mM KH_2_PO_4_ pH 7.5) supplemented with 1% Triton, 10 µg/mL DNaseI, 5 mM MgCl2, lysosome, and cOmplete™ EDTA-free Protease Inhibitor Cocktail (Roche)) and was incubated for 1h on ice. Remaining DNA was destroyed by sonicated, and the cell debris was pelleted by centrifugation (1h, 16,000 x g). The GST-tagged protein was purified on GSH Sepharose 4 Fast Flow Media (Cytiva) by allowed to associate for 1 h at 4°C with the beads. The unbound proteins were washed away with 30 column volumes PBS. The protein was cleaved on the beads with thrombin in PBS for 18 h at 4°C. To remove thrombin from the cleaved protein, the protein was applied on a HiTrap Benzamidine FF (Cytiva), which was equilibrated and eluted cleaved protein with binding buffer (50mM Tris HCl pH7.4 500mM NaCl). The pure protein was dialyzed to H2O using SnakeSkin Dialysis Tubing 10,000 MWCO. The protein concentration was determined spectrometric and the purity was confirmed through SDS-PAGE.

### NMR titration

NMR titration experiments were performed on a 600 MHz Buker Avance NEO spectrometer equipped with a 5 mm TCI cryogenic probe at 25 °C on a sample of 200 μM ^15^N-labelled-Hsp90-NTD in an unbuffered solution of 30% D_2_O in H_2_O. ^1^H,^15^N-HSQC spectra were recorded using solvent suppression, 1024 points in f1, 256 increments in f1, 32 transients, and a relaxation delay of 0.2s. Spectra were recorded at 0, 0.25, 0.50, 1.00, 1.50, 2.50 and 5.00 molar equivalents of *LLN-containing peptide*. No precipitation had occurred after addition of 5.00 equivalents. Crosspeaks of the bound and unbound protein were present, indicative of slow chemical exchange. Chemical shift perturbations (CSPs, Supplementary Table 7) were calculated as the average chemical shift changes observed for ^1^H and ^15^N nuclei (Δδ_avg_) between the spectra of 0 and 5.00 equivalents of *LLN-containing peptide* using equation S1.

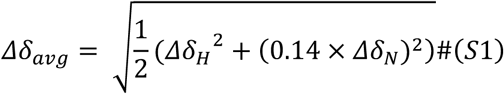

In equation S1, Δδ_avg_ is the averaged chemical shift perturbation, Δδ_H_ and Δδ_N_ are the chemical shift perturbations in the ^1^H and ^15^N dimensions respectively. A scaling factor of 0.14 was applied to the nitrogen chemical shift perturbation to account for differences in the chemical shift ranges.

^1^H,^15^N-HSQC spectra of ^15^N-labelled-HSP90-NTD were acquired at increasing concentrations of *LLN-containing peptide* PQLRSLLLNPPPPQTY-CONH2, up to 5 molar equivalents, in order to elucidate the protein binding site of the peptide. Signals of free and bound HSP90 were observed during the titration, indicating slow exchange between the free and bound forms. Residues with chemical shift perturbation (CSP) values greater than 2 standard deviations were considered significant and indicative of direct binding.

### LUMIER assay

The LUMIER assays were adapted from a previously published protocol^12^. HEK293T cells stably expressing HSP90AB1 fused to Nanoluc at the N-terminus were seeded into clear 96-well plates at 25,000 – 30,000 cells per well. The following day, each well was transfected with 75 ng of 3xFLAG-V5 tagged bait proteins and 75 ng of either EGFP fused to an HSP90 peptide or EGFP on its own using polyethylenimine (Polysciences, 24765). Two days after transfection, cells were washed with 1xPBS using an automated plate washer (Biotek) and then lysed with ice-cold HENG buffer (20 mM HEPES pH 7.9, 150 mM NaCl, 20 mM Na_2_MoO_4_, 2 mM EDTA pH 8, 0.5% Triton X-100, 5% glycerol) containing protease inhibitors (1 μg/ml aprotinin, 1 μg/ml leupeptin, 1 μg/ml pepstatin, 0.2 mM PMSF). Alternatively, as a positive control for HSP90 inhibition, cells were instead transfected with 150 ng of 3xFLAG-V5 tagged bait proteins and treated with 1 µM of Ganetespib (STA-9090) for 1 hour before the cell wash and lysis. Lysates were then transferred into opaque, white, high binding 384-well plates (Greiner Bio-One) that were pre-coated with monoclonal anti-FLAG M2 antibody (Sigma-Aldrich, F1804) and blocked (1% BSA, 5% sucrose, 0.5% Tween-20, 1xPBS). The 384-well plates were incubated for 3 hours at 4°C with mild shaking and then washed with HENG buffer without protease inhibitors using an automated plate washer. Luminescence was measured with a BioTek Synergy Neo microplate reader five minutes after adding furimazine luciferase reagent dissolved 1:200 in luciferase buffer (20 mM Tris-HCl pH 7.5, 1 mM EDTA, 150 mM KCl, 0.5% Tergitol NP9). The Luciferase buffer was then removed from each well and HRP-conjugated anti-FLAG antibody (1:10,000, Sigma-Aldrich, A8592) diluted in ELISA buffer (1× PBS, 1% goat serum, 1% Tween 20) was added to wells. plates were incubated for 90 minutes at room temperature with mild shaking and then washed with 0.1% Tween-20 in 1xPBS using an automated plate washer. ELISA signal was measured using a BioTek Synergy Neo microplate reader one minute after adding SuperSignal ELISA Pico Chemiluminescent substrate (ThermoFisher Scientific, 37069).

ELISA values measuring the expression of 3xFLAG-V5 tagged bait proteins in each well were used to remove any proteins not expressed above background levels. ELISA values were also used to adjust luminescence values in control conditions (EGFP transfection or DMSO) based on the expression of bait proteins relative to the HSP90 peptide or HSP90 inhibitor conditions. Final LUMIER interactions scores were then derived by normalizing the luminescence values in each well to the average luminescence of non-interacting negative controls (e.g., 3xFLAG-tagged EGFP) and empty wells in the same 384-well plate.

