## Supplementary Table 7 for "An Atlas of Short Linear Motif-Mediated Human Protein-Protein Interactions"

**Table S7.** Chemical shift perturbations (CSP) observed for Hsp90-NTD with 5 equivalents of LLN-containing peptide. Only residues for which CSP values could be determined are given. Residues with a CSP greater than 1 and 2 standard deviation calculated (σ = 0.0364) are indicated by an x.

|  | Before | | After | |  |  |  |
| --- | --- | --- | --- | --- | --- | --- | --- |
| Residue | δH (ppm) | δN (ppm) | δH (ppm) | δN (ppm) | CSP | > 1σ | > 2σ |
| **18E** | 8.44 | 126.47 | 8.44 | 126.47 | 0.001 |  |  |
| **20F** | 8.66 | 125.32 | 8.66 | 125.32 | 0.001 |  |  |
| **21A** | 8.24 | 122.56 | 8.24 | 122.51 | 0.004 |  |  |
| **22F** | 7.66 | 118.61 | 7.66 | 118.70 | 0.009 |  |  |
| **25G** | 7.64 | 106.10 | 7.64 | 106.10 | 0.001 |  |  |
| **26I** | 7.26 | 120.14 | 7.25 | 120.08 | 0.008 |  |  |
| **30M** | 8.32 | 116.98 | 8.32 | 116.94 | 0.005 |  |  |
| **31S** | 7.59 | 111.94 | 7.61 | 111.89 | 0.015 |  |  |
| **32L** | 7.80 | 122.08 | 7.81 | 122.07 | 0.007 |  |  |
| **34I** | 8.08 | 116.68 | 8.07 | 116.63 | 0.006 |  |  |
| **35N** | 7.67 | 114.70 | 7.69 | 114.70 | 0.011 |  |  |
| **36T** | 7.29 | 116.69 | 7.29 | 116.85 | 0.017 |  |  |
| **41K** | 7.88 | 117.33 | 7.83 | 117.07 | 0.044 | x |  |
| **42E** | 8.35 | 114.57 | 8.37 | 114.44 | 0.021 |  |  |
| **43I** | 6.83 | 112.46 | 6.86 | 112.55 | 0.022 |  |  |
| **46R** | 6.43 | 117.51 | 6.25 | 117.40 | 0.128 | x | x |
| **50S** | 8.24 | 117.07 | 8.23 | 116.85 | 0.022 |  |  |
| **53S** | 8.54 | 116.17 | 8.51 | 115.93 | 0.029 |  |  |
| **54D** | 8.15 | 119.66 | 8.22 | 119.66 | 0.053 | x |  |
| **56L** | 8.44 | 124.67 | 8.42 | 124.75 | 0.013 |  |  |
| **57D** | 8.79 | 120.01 | 8.83 | 120.06 | 0.025 |  |  |
| **58K** | 7.75 | 117.47 | 7.73 | 117.42 | 0.013 |  |  |
| **61Y** | 8.66 | 118.52 | 8.66 | 118.61 | 0.01 |  |  |
| **64L** | 7.35 | 121.81 | 7.34 | 121.81 | 0.002 |  |  |
| **65T** | 6.99 | 106.80 | 6.99 | 106.75 | 0.006 |  |  |
| **66D** | 7.30 | 118.08 | 7.29 | 118.04 | 0.006 |  |  |
| **68S** | 8.05 | 115.40 | 8.04 | 115.34 | 0.01 |  |  |
| **70L** | 7.17 | 110.88 | 7.16 | 110.84 | 0.006 |  |  |
| **72S** | 7.44 | 109.26 | 7.42 | 109.12 | 0.015 |  |  |
| **73G** | 7.64 | 111.45 | 7.63 | 111.45 | 0.011 |  |  |
| **75E** | 7.62 | 116.90 | 7.59 | 116.76 | 0.02 |  |  |
| **77H** | 7.63 | 118.96 | 7.61 | 119.31 | 0.038 | x |  |
| **78I** | 8.10 | 117.77 | 8.13 | 118.17 | 0.044 | x |  |
| **79N** | 9.69 | 125.32 | 9.83 | 125.32 | 0.103 | x | x |
| **80L** | 8.97 | 122.25 | 9.23 | 121.68 | 0.19 | x | x |
| **81I** | 9.31 | 120.23 | 9.30 | 119.97 | 0.027 |  |  |
| **83N** | 9.11 | 121.55 | 9.39 | 121.59 | 0.199 | x | x |
| **84K** | 9.39 | 124.97 | 9.32 | 124.67 | 0.063 | x |  |
| **86D** | 7.58 | 116.50 | 7.56 | 116.41 | 0.018 |  |  |
| **88T** | 8.02 | 106.14 | 8.01 | 106.05 | 0.01 |  |  |
| **90T** | 7.99 | 124.23 | 7.86 | 124.40 | 0.092 | x | x |
| **91I** | 9.41 | 127.96 | 9.44 | 128.00 | 0.017 |  |  |
| **92V** | 9.60 | 129.85 | 9.68 | 129.80 | 0.06 | x |  |
| **93D** | 9.44 | 123.52 | 9.45 | 123.39 | 0.015 |  |  |
| **94T** | 7.16 | 109.56 | 7.16 | 109.56 | 0.002 |  |  |
| **95G** | 9.52 | 108.90 | 9.55 | 108.95 | 0.015 |  |  |
| **96I** | 6.82 | 115.80 | 6.82 | 115.71 | 0.01 |  |  |
| **97G** | 7.64 | 103.68 | 7.62 | 103.57 | 0.017 |  |  |
| **99T** | 8.01 | 112.24 | 8.02 | 112.33 | 0.009 |  |  |
| **101A** | 7.87 | 116.68 | 7.87 | 116.68 | 0 |  |  |
| **102D** | 7.39 | 117.60 | 7.39 | 117.60 | 0.003 |  |  |
| **104I** | 7.56 | 116.15 | 7.56 | 116.41 | 0.026 |  |  |
| **105N** | 8.28 | 117.60 | 8.28 | 117.60 | 0.003 |  |  |
| **106N** | 8.75 | 116.28 | 8.77 | 116.24 | 0.01 |  |  |
| **108G** | 7.62 | 105.66 | 7.63 | 105.70 | 0.01 |  |  |
| **119M** | 7.84 | 117.47 | 7.85 | 117.51 | 0.01 |  |  |
| **121A** | 7.76 | 124.18 | 7.75 | 124.14 | 0.005 |  |  |
| **124A** | 7.37 | 120.32 | 7.37 | 120.32 | 0.001 |  |  |
| **129S** | 8.51 | 114.48 | 8.51 | 114.44 | 0.004 |  |  |
| **134F** | 7.32 | 115.62 | 7.30 | 115.58 | 0.014 |  |  |
| **136V** | 6.56 | 107.94 | 6.57 | 107.94 | 0.004 |  |  |
| **137G** | 8.71 | 107.72 | 8.72 | 107.72 | 0.005 |  |  |
| **138F** | 9.10 | 123.52 | 9.11 | 123.48 | 0.007 |  |  |
| **140S** | 7.95 | 114.30 | 7.95 | 114.35 | 0.006 |  |  |
| **144V** | 6.58 | 102.50 | 6.57 | 102.45 | 0.008 |  |  |
| **145A** | 7.63 | 124.31 | 7.63 | 124.27 | 0.004 |  |  |
| **146E** | 8.50 | 118.08 | 8.51 | 118.17 | 0.011 |  |  |
| **148V** | 8.02 | 125.90 | 8.01 | 125.76 | 0.015 |  |  |
| **149T** | 9.02 | 124.93 | 9.03 | 124.97 | 0.006 |  |  |
| **150V** | 10.45 | 128.44 | 10.44 | 128.40 | 0.004 |  |  |
| **151I** | 9.58 | 128.75 | 9.62 | 128.81 | 0.027 |  |  |
| **152T** | 9.14 | 122.16 | 9.15 | 122.12 | 0.01 |  |  |
| **154H** | 9.68 | 131.95 | 9.65 | 132.00 | 0.018 |  |  |
| **155N** | 9.58 | 126.03 | 9.57 | 126.03 | 0.011 |  |  |
| **160Y** | 8.64 | 124.45 | 8.62 | 124.14 | 0.034 |  |  |
| **161A** | 9.28 | 120.06 | 9.30 | 119.97 | 0.014 |  |  |
| **162W** | 10.31 | 130.13 | 10.32 | 130.15 | 0.007 |  |  |
| **164S** | 8.20 | 113.38 | 8.19 | 113.34 | 0.009 |  |  |
| **166A** | 9.06 | 119.66 | 9.07 | 119.66 | 0.006 |  |  |
| **167G** | 8.43 | 106.71 | 8.44 | 106.71 | 0.006 |  |  |
| **168G** | 8.45 | 106.36 | 8.45 | 106.36 | 0.006 |  |  |
| **169S** | 7.76 | 115.53 | 7.76 | 115.53 | 0.001 |  |  |
| **170F** | 8.78 | 119.66 | 8.78 | 119.62 | 0.004 |  |  |
| **171T** | 8.73 | 109.48 | 8.74 | 109.48 | 0.002 |  |  |
| **172V** | 8.82 | 118.17 | 8.83 | 118.12 | 0.005 |  |  |
| **173R** | 9.07 | 124.14 | 9.08 | 124.18 | 0.007 |  |  |
| **174T** | 9.05 | 118.52 | 9.06 | 118.52 | 0 |  |  |
| **176T** | 7.98 | 115.71 | 7.98 | 115.71 | 0.004 |  |  |
| **177G** | 7.78 | 110.13 | 7.77 | 110.09 | 0.005 |  |  |
| **180M** | 9.40 | 123.96 | 9.40 | 123.92 | 0.007 |  |  |
| **181G** | 8.59 | 111.45 | 8.60 | 111.50 | 0.012 |  |  |
| **182R** | 7.41 | 120.06 | 7.42 | 120.14 | 0.011 |  |  |
| **183G** | 9.15 | 115.53 | 9.16 | 115.58 | 0.011 |  |  |
| **184T** | 7.71 | 115.89 | 7.73 | 115.93 | 0.011 |  |  |
| **185K** | 10.01 | 128.40 | 10.02 | 128.75 | 0.036 |  |  |
| **186V** | 9.15 | 126.51 | 9.21 | 126.77 | 0.051 | x |  |
| **187I** | 9.94 | 127.87 | 9.92 | 127.74 | 0.016 |  |  |
| **188L** | 8.86 | 125.90 | 8.85 | 125.85 | 0.004 |  |  |
| **189H** | 8.29 | 124.84 | 8.32 | 124.97 | 0.028 |  |  |
| **192E** | 8.98 | 120.63 | 8.99 | 120.63 | 0.008 |  |  |
| **193D** | 8.35 | 113.21 | 8.36 | 113.21 | 0.006 |  |  |
| **195T** | 7.39 | 106.45 | 7.41 | 106.45 | 0.011 |  |  |
| **196E** | 8.82 | 123.92 | 8.82 | 123.83 | 0.01 |  |  |
| **197Y** | 6.62 | 114.70 | 6.61 | 114.70 | 0.011 |  |  |
| **199E** | 7.29 | 116.68 | 7.31 | 115.97 | 0.072 | x |  |
| **200E** | 9.15 | 126.51 | 9.21 | 126.77 | 0.051 | x |  |
| **201R** | 8.89 | 116.24 | 8.91 | 116.02 | 0.029 |  |  |
| **202R** | 6.75 | 119.49 | 6.70 | 119.22 | 0.045 | x |  |
| **203I** | 8.10 | 117.77 | 8.13 | 118.17 | 0.044 | x |  |
| **204K** | 8.41 | 116.81 | 8.43 | 117.11 | 0.034 |  |  |
| **205E** | 7.66 | 118.61 | 7.66 | 118.70 | 0.009 |  |  |
| **209K** | 7.48 | 116.41 | 7.47 | 116.06 | 0.035 |  |  |
| **210H** | 7.78 | 113.43 | 7.90 | 112.86 | 0.098 | x | x |
| **211S** | 8.13 | 115.45 | 8.26 | 116.32 | 0.124 | x | x |
| **213F** | 8.13 | 115.45 | 8.26 | 116.32 | 0.124 | x | x |
| **214I** | 7.10 | 121.59 | 7.10 | 123.00 | 0.139 | x | x |
| **215G** | 9.15 | 115.53 | 9.24 | 116.10 | 0.088 | x | x |
| **216Y** | 6.71 | 117.69 | 6.77 | 118.26 | 0.069 | x |  |
